# Iron–endoplasmic reticulum–extracellular matrix axis regulates cancer cell invasion

**DOI:** 10.64898/2026.06.29.735430

**Authors:** Arun Asif, Kavya Panjwani, Kavya Nair, Percy Smith, Oruko Dancan, Isaiah Crosbourne, Joseph DeLuca, Taylor Humphrey, Ramon Bossardi Ramos, David T. Corr, Teresita Padilla-Benavides, Margarida Barroso

## Abstract

Intracellular iron homeostasis is increasingly recognized as a regulator of cancer cell behavior, but how iron distribution influences extracellular matrix (ECM) organization and invasion remains poorly understood. Here, we show that loss of divalent metal transporter 1 (DMT1/SLC11A2) disrupts intracellular iron homeostasis and promotes cancer cell invasion through an iron–ER–ECM axis. In MDA-MB-231 cells, DMT1 knockout (KO) reduced total iron content but increased the labile iron pool (LIP) in both 2D and 3D culture models, indicating altered intracellular iron distribution. Across transcriptomic and phenotypic readouts, DMT1-dependent effects were more evident in 3D than in 2D models, with DMT1 KO inducing endoplasmic reticulum (ER) stress and impaired collagen/ECM organization. Functionally, the DMT1-loss phenotype was marked by reduced 2D motility, whereas in 3D spheroid models DMT1 KO cells displayed enhanced invasive outgrowth in both Matrigel and collagen matrices. Iron chelation further modulated this phenotype in a DMT1-dependent manner. Pharmacologic induction of ER stress phenocopied the loose spheroid architecture and invasive behavior, supporting ER stress as a mechanistic link between altered iron handling and ECM destabilization. Together, these findings identify intracellular iron distribution, rather than total iron abundance alone, as a determinant of ECM integrity and context-dependent cancer cell invasion.

**One Sentence Summary:** Iron misrouting, not iron excess, drives ER stress and collagen failure that unleash invasion in triple-negative breast cancer.

## INTRODUCTION

Iron is essential for cancer cell proliferation, mitochondrial metabolism, and redox homeostasis, and many cancer cells acquire an iron-accumulating phenotype to support these demands (*1–3*). At the same time, excess iron can promote reactive oxygen species (ROS) generation through Fenton chemistry (Fe²⁺ + H₂O₂ → Fe³⁺ + •OH + OH⁻) (*4*), thereby perturbing protein folding and other iron-sensitive cellular processes (*5, 6*). Although most studies have focused on total iron content as a driver of iron-dependent cancer cell behavior, these processes may also be shaped by how iron is distributed within the cell. This distinction may be especially important for cancer cell invasion, because iron-dependent responses are strongly influenced by cellular context (*7*). Conventional 2D monolayer cultures capture cell spreading and migration on planar surfaces but do not reproduce the spatial constraints, multicellular organization, matrix interactions, and metabolic gradients present in tumors. In contrast, three-dimensional (3D) tumor spheroids better model these features and may therefore reveal invasion-related phenotypes that are not apparent in 2D systems (*8–11*). This issue is particularly relevant to divalent metal transporter 1 (*DMT1*, also known as *SLC11A2* or NRAMP2), a proton-coupled metal ion transporter that mediates cellular ferrous iron uptake and endosomal iron release during transferrin-dependent trafficking and recycling (*12, 13*).

Previous work established active transferrin-dependent iron uptake, trafficking and recycling in MDA-MB-231 cells and spheroids (*14, 15*). Label-free Raman imaging directly detected intracellular iron-bound transferrin and revealed delayed endosomal iron release in MDA-MB-231 cells (*14*). Iron-loaded transferrin was also internalized throughout 3D spheroids, with greater spatial heterogeneity in larger aggregates (*16*). DMT1 localizes to transferrin-containing endosomes and promotes endosome–mitochondria interactions and mitochondrial transfer of transferrin-derived iron (*17, 18*); DMT1 loss impairs this process, whereas re-expression restores it (*17*). DMT1 is also upregulated in multiple cancers and has been linked to iron accumulation, ROS generation, metabolic reprogramming, ferroptosis, and EMT (*17, 19–22*). Importantly, DMT1 depletion increased the labile iron pool and metastatic outgrowth in both human MDA-MB-231 xenografts and murine E0771 TNBC cells in a species-matched syngeneic model (*17*). Together, these findings support a DMT1-dependent defect in intracellular iron routing rather than impaired transferrin uptake and led us to test whether altered iron homeostasis regulates invasion in a context-dependent manner, with reduced motility in 2D diverging from invasive behavior in 3D architectures that depend on ECM remodeling, multicellular adaptation, and structural containment.

One process likely to be especially sensitive to such context-dependent iron handling is extracellular matrix (ECM) organization. ECM architecture is a central regulator of tumor structure and cancer cell invasion (*23*), and collagen is particularly important because its loss or disorganization weakens matrix integrity and facilitates invasive behavior (*24*). Collagen synthesis and maturation depend on iron-dependent enzymatic reactions together with endoplasmic reticulum (ER)-localized oxidative folding processes, suggesting that altered intracellular iron homeostasis may disrupt ECM homeostasis (*25, 26*). Disruption of ER proteostasis activates the unfolded protein response (UPR), an ER stress-signaling program that helps restore protein-folding capacity but can also impair secretory and matrix-producing functions when chronically engaged (*27, 28*). We therefore asked whether the phenotype caused by *DMT1* loss reflects defects in ECM maintenance that become apparent under 3D tumor-like conditions. Using previously generated MDA-MB-231 cell lines with genetically modified *DMT1* expression, including the *DMT1* KO and the stable *DMT1*-GFP overexpression (OE) cells, we compared their phenotypes in 2D monolayers and liquid-overlay 3D tumor spheroids (*17*). We show that loss of DMT1 disrupts intracellular iron distribution, induces ER stress, impairs collagen production, weakens ECM architecture, and promotes cancer cell invasion in 3D tumor models. These findings identify iron homeostasis as a regulator of ECM integrity and support *DMT1* as a suppressor of cancer cell invasive behavior.

## RESULTS

### Spheroid size alters the transcriptional baseline of MDA-MB-231 3D spheroids

Defining how cell culture dimensionality and spheroid size affect baseline gene expression is important for interpreting DMT1*-*dependent phenotypes in 3D systems, particularly because iron metabolism intersects with hypoxia signaling (HIF pathways) and oxidative metabolism (*29*). We therefore first characterized the transcriptional landscape of wild-type (WT) MDA-MB-231 cells grown as two-dimensional (2D) monolayers, liquid-overlay small (LOS) spheroids (8,000 cells/spheroid, approximately 300 to 500 µm in diameter), or liquid-overlay large (LOL) spheroids (25,000 cells/per spheroid, approximately 500 to 800 µm in diameter; **Fig. 1A** and **table S2**). Heatmap analysis revealed clear transcriptional differences between 2D and 3D cultures, with the most prominent changes involving cell-cycle, mitochondrial, and hypoxia-related programs. Cell cycle genes including *CCNA1*, *MKI67*, and *BUB1* were significantly reduced in 3D cultures, with a more pronounced decrease in LOL spheroids (**Fig. 1A, left**). Consistent with dormancy in spheroid cores (*30, 31*), mitochondrial genes such as *TOMM34*, *MRPL13*, and *ATP5B* were also downregulated in both 3D models, indicating reduced oxidative metabolism (**Fig. 1A, middle**). In contrast, hypoxia-inducible genes including *HIF1A*, *VEGFA*, *PLOD2*, and *EGLN1* were elevated in LOL compared with 2D and LOS, consistent with greater hypoxic stress in larger spheroids (**Fig. 1A, right**).

**Fig. 1.**
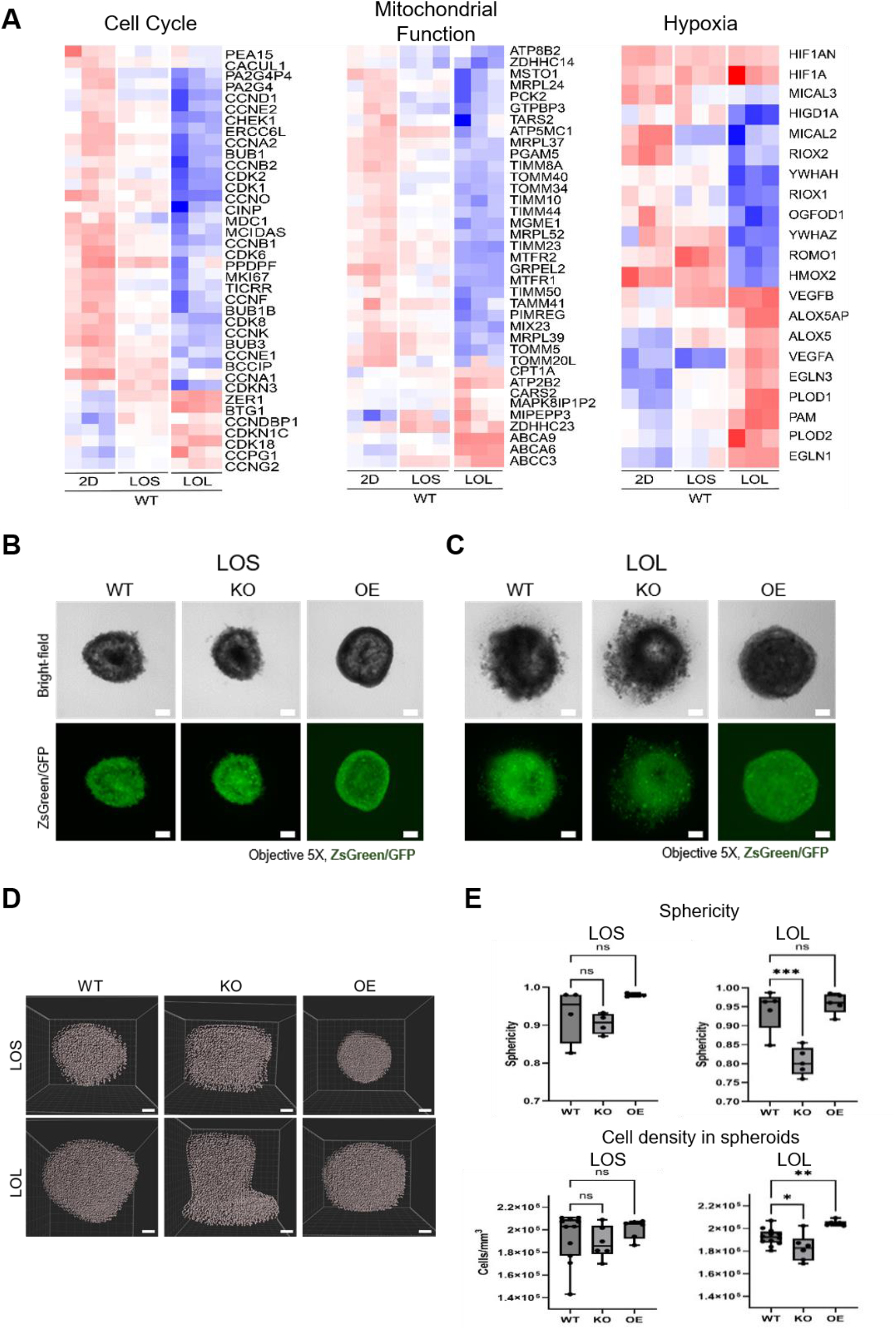
*DMT1* regulates spheroid compaction and spheroid size shapes baseline transcriptional programs in MDA-MB-231 3D models. (A) Heatmaps showing expression of representative cell-cycle, mitochondrial, and hypoxia-related genes in WT MDA-MB-231 cells grown in 2D monolayer culture, liquid-overlay small (LOS), and liquid-overlay large (LOL) spheroids. The full list of differentially expressed genes underlying these heatmaps is provided in **data file S1**. (B-C) Representative brightfield images (top row) and fluorescence images (bottom row) of LOS (B) and LOL (C) spheroids formed by WT, *DMT1* KO, and *DMT1*-GFP OE MDA-MB-231 cells after 4 days in culture. WT and KO cells were visualized with ZsGreen and OE cells with DMT1-GFP. Scale bar = 200 µm. (D) Representative OCT-based 3D reconstructions of LOS and LOL spheroids from WT, KO, and OE cells. (E) Quantitative OCT analysis of spheroid sphericity and cell density in LOS and LOL (n = 6). *DMT1* KO spheroids showed reduced sphericity and cell density, whereas *DMT1* OE spheroids showed increased sphericity and density, with the strongest differences observed in LOL spheroids. Data represent mean ± SEM (n = 6); one-way ANOVA (GraphPad Prism). *p < 0.05, **p < 0.01, ***p < 0.001; ns, not significant

Because *DMT1* KO transcriptomic profiles were reported previously (*17*), the present analysis focused on establishing the 2D *vs.* 3D transcriptional baseline and determining whether *DMT1* overexpression alters that baseline program. The DMT1-GFP OE construct was originally generated and characterized by Barra et al. (*17*), including validation of its expression and subcellular localization. Stable overexpression of DMT1-GFP MDA-MB-231 cells (OE) was independently validated by western blot (**fig. S1**). RNA sequencing analysis was performed on 2D, LOS, and LOL cultures of stable MDA-MB-231 OE cells. Principal component analysis (PCA) showed that spheroid size, rather than *DMT1* expression, was the dominant source of transcriptional variance **(fig. S2).** PC1 (33% of variance) separated 2D cultures from 3D spheroids, with LOL models showing the largest displacement in both WT and OE cells, whereas 2D and LOS conditions clustered more closely. PC2 (23% of variance) showed minor separation between WT and OE within each condition. Consistent with this, the largest transcriptional differences in both WT and OE cells were observed in the 2D *vs.* LOL comparison, whereas LOL *vs.* LOS showed relatively few changes (**figs. S3 and S4; complete differentially expressed gene lists in data files S1 and S3**). Together, these findings indicate that dimensionality and spheroid size define the dominant transcriptional baseline of the model. LOL spheroids showed the strongest hypoxia-associated and metabolic divergence from 2D culture and were selected for interrogating DMT1-dependent effects on tumor architecture and cancer cell invasion.

### *DMT1* promotes spheroid compaction in larger tumor spheroids

We next asked whether DMT1 affects spheroid structural integrity in LOL spheroids. WT, *DMT1* KO and OE MDA-MB-231 cells were grown as LOS and LOL spheroids for 4 days. WT and KO cells were labeled with ZsGreen for visualization whereas OE cells expressed DMT1-GFP. Brightfield and fluorescence imaging showed that WT and *DMT1* OE cells formed compact, spherical spheroids, whereas *DMT1* KO cells formed looser aggregates with reduced sphericity, an effect that was more pronounced in LOL spheroids (**Fig. 1B-D**). To quantify these 3D morphological differences, we used optical coherence tomography (OCT), which enables non-destructive structural analysis of intact spheroids (*16, 32*). OCT confirmed marked genotype-dependent differences in 3D morphology, particularly in LOL spheroids (**Fig. 1C-D**). *DMT1* OE LOL spheroids were the most spherical, approaching a sphericity value of 1, consistent with highly rounded, smooth-edged aggregates, whereas *DMT1* KO LOL spheroids showed significantly reduced sphericity relative to WT and OE (**Fig. 1E**). While spheroid volume remained comparable across all cell types, *DMT1* KO LOL spheroids exhibited significantly lower cell density, consistent with their loose aggregation phenotype. In contrast, *DMT1* OE increased spheroid compactness and density, indicating that *DMT1* expression promotes structural cohesion in 3D culture. In particular, the *DMT1* KO and OE genotype-dependent differences were more evident in LOL than in LOS spheroids, where the effects were not significant (**Fig. 1E**). Two-photon imaging further supported these structural differences. OE spheroids displayed a densely packed core that restricted DAPI penetration despite uniform ZsGreen fluorescence, indicating an intact and highly compact cellular architecture (**fig. S5**). In contrast, KO spheroids showed more permissive dye penetration and looser internal organization. These structural differences suggest that DMT1 promotes compaction and structural integrity in 3D constructs, prompting us to examine whether these differences are associated with altered iron homeostasis.

### *DMT1* depletion increases labile iron despite reduced total iron content

The altered spheroid morphology suggests that *DMT1* supports the structural cohesion of multicellular tumor aggregates through regulation of iron homeostasis. We therefore examined total and LIP levels in WT, *DMT1* KO and OE MDA-MB-231 cells. Total cellular iron measurements using atomic absorption spectroscopy (AAS) showed that *DMT1* KO cells contained significantly less iron than WT and *DMT1* OE cells under both 2D and LOL culture conditions (**Fig. 2A**) (*33, 34*). Counterintuitively, *DMT1* KO cells exhibited increased LIP levels, despite their reduced total iron content. Using FerroOrange, a fluorescent dye that selectively detects labile Fe^2+^, we observed significantly increased fluorescence intensity levels in *DMT1* KO cells compared to WT and *DMT1* OE cells, under 2D and LOL conditions (**Fig. 2B**). Quantitative analysis confirmed that *DMT1* KO cells showed approximately 2-fold higher LIP levels than WT, consistent with our previous findings (*17*) (**Fig. 2B, right panel**). This combination of reduced total iron and increased LIP is inconsistent with a simple generalized iron-deficiency state and instead indicates impaired intracellular iron distribution.

**Fig. 2.**
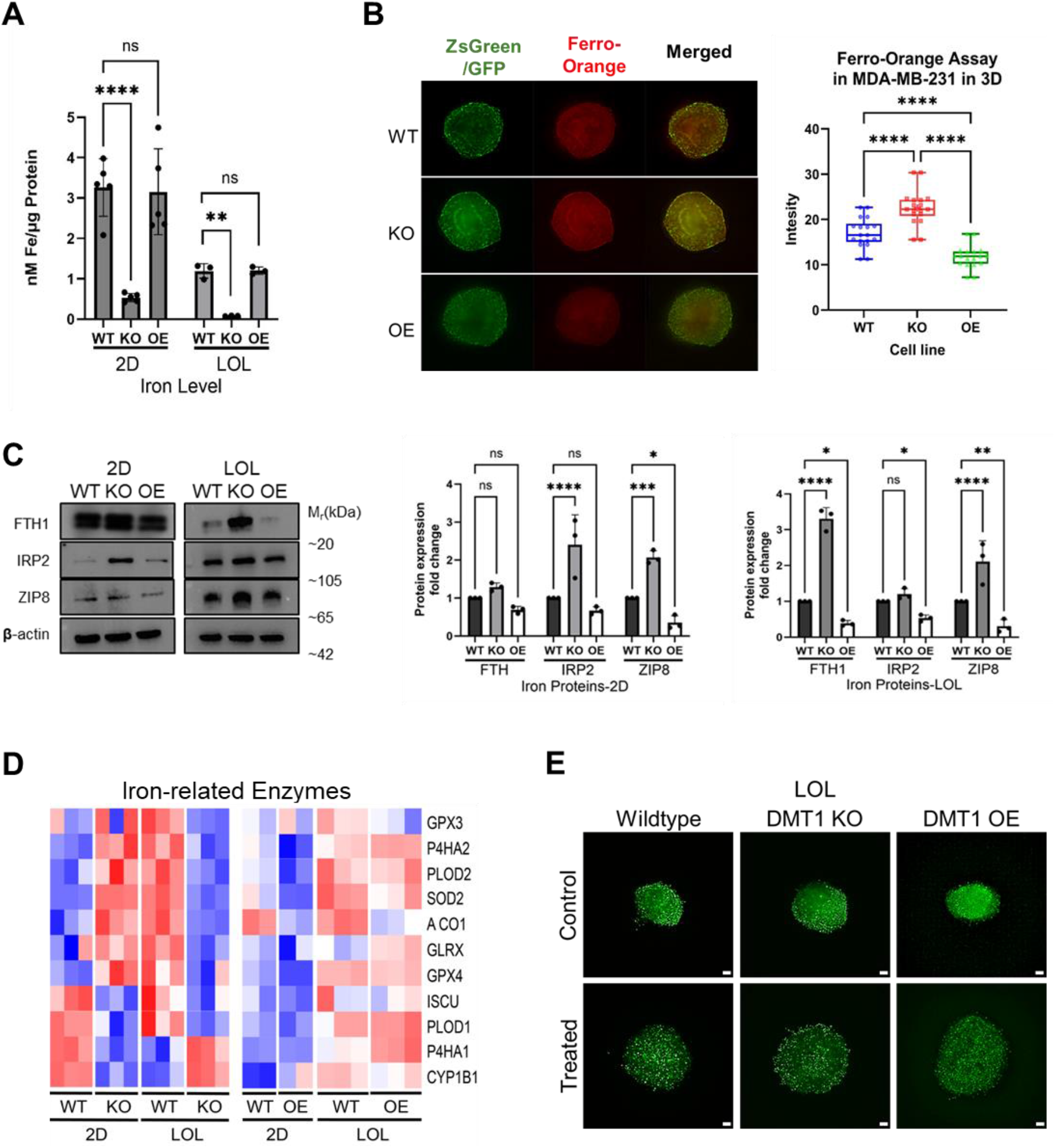
*DMT1* loss increases labile iron despite reduced total iron content. **(A)** Total iron levels (nmol/µg protein) in WT, *DMT1* KO and OE MDA-MB-231 cells grown under 2D and LOL culture conditions. **(B)** Representative FerroOrange images of LOL spheroids showing ZsGreen/GFP fluorescence, FerroOrange signal, and merged images for WT, KO, and OE cells (left and middle panels). Quantification of Ferro-Orange intensity in MDA-MB-231 cells under 2D monolayers and LOL spheroids (right panel), indicating increased LIP buildup in *DMT1* KO cells compared to WT and OE (***p<0.001) in 2D and 3D conditions. **(C)** Representative western blots (left) and quantification (right) of iron-regulatory proteins (FTH1, IRP2, ZIP8) in WT, KO, and OE cells under 2D and LOL conditions, with β-actin as loading control. **(D)** Heatmap showing relative expression of representative iron-related enzymes in WT, KO, and OE cells under 2D and LOL conditions. Values are normalized by gene to visualize relative up-or downregulation across conditions. The merged differential expression dataset used to generate these heatmaps is provided in **data file S2**. **(E)** Representative LOL spheroids formed under iron-chelated conditions with deferoxamine (DFO). WT and KO spheroids remained compact, whereas OE spheroids formed looser, less cohesive aggregates under iron-restricted conditions; Scale bar = 200 µm. FerroOrange intensity was quantified from 6 spheroids per condition across two independent experiments, with ≥10 images per spheroid and 4-5 ROIs per image; values from all images and ROIs were averaged per spheroid, so that each spheroid represented one biological replicate (n = 6). Western blots represent 5 independent repeats per protein. Data represent mean ± SEM; one-way ANOVA (GraphPad Prism). *p < 0.05, **p < 0.01, ***p < 0.001, ****p < 0.0001; ns, not significant.

Western blot analysis of iron-regulatory proteins further supported a noncanonical iron-redistribution state in DMT1 KO cells. Ferritin heavy chain (FTH1) and iron regulatory protein 2 (IRP2) were significantly increased in *DMT1* KO cells, under both 2D and LOL systems **(Fig. 2C)**. In addition, ZIP8 (SLC39A8), a transporter known to mobilize iron, zinc and manganese (*35*), was also elevated in *DMT1* KO cells, suggesting a compensatory response to altered iron handling. Consistent with this, transcriptomic analysis showed downregulation of representative iron-related enzymes in *DMT1* KO cells, particularly in LOL spheroids (**Fig. 2D**). To test whether spheroid structure depends on iron availability, we next treated LOL spheroids with the iron chelator deferoxamine (DFO). DFO altered spheroid aggregation in a genotype-dependent manner: WT and *DMT1* KO spheroids largely maintained compact structures under iron-restricted conditions, whereas *DMT1* OE spheroids failed to fully aggregate and instead formed loose, less cohesive spheroids, indicating that OE spheroid compaction is more sensitive to iron restriction (**Fig. 2E**). Together, these findings show that spheroid architecture responds to iron restriction in a DMT1-dependent manner and prompted us to examine ECM organization and collagen synthesis.

### Loss of *DMT1* impairs collagen production and ECM organization

Because spheroid cohesion depends in part on ECM deposition, we asked whether the loose aggregation of DMT1 KO spheroids reflects impaired ECM production. We examined the expression of ER-resident proteins involved in collagen biosynthesis and maturation, such as protein disulfide isomerase (PDI; disulfide bond formation), prolyl-4-hydroxylase 1 (P4HA1; proline hydroxylation), ER oxidoreductase (ERO1; PDI activator), and COL1A1. *DMT1* KO cells showed elevated PDI and P4HA1 levels, but COL1A1 expression was significantly decreased in both 2D and LOL conditions, with a more pronounced reduction in LOL spheroids (**Fig. 3A**). ERO1 was also reduced in KO cells, despite elevated PDI and P4HA1, suggesting that *DMT1* loss disrupts ER oxidative folding and collagen maturation not directly through reduced iron abundance, but through impaired coordination of iron-dependent biosynthesis and ER proteostasis. Because P4HA1 requires iron as a cofactor, altered intracellular iron partitioning may further limit functional collagen-processing capacity despite elevated enzyme expression. In contrast, *DMT1* OE cells showed higher ERO1 protein expression, especially in LOL spheroids where levels were significantly higher than WT (**Fig. 3A**).

**Fig. 3.**
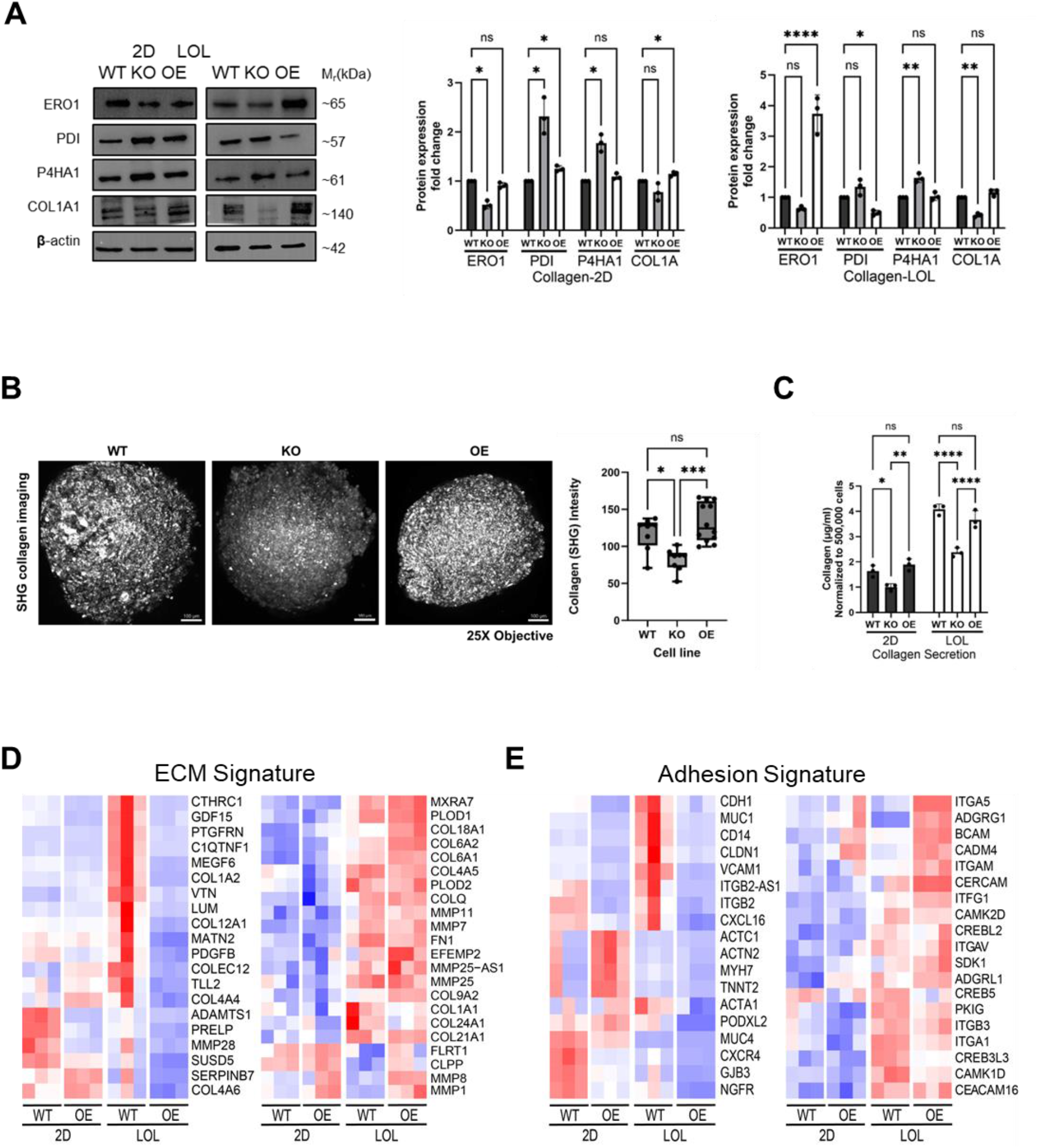
*DMT1* loss impairs collagen production and ECM organization. **(A)** Representative western blot images (left) and quantification (right) of collagen biosynthesis and maturation markers (ERO1, PDI, P4HA1 and COL1A1) in WT and *DMT1* KO and OE MDA-MB-231 cells under 2D and LOL conditions, with β-actin as loading control (left). **(B)** Representative second harmonic generation (SHG) images of collagen in WT, KO, and OE LOL spheroids. SHG imaging was performed at 820 nm excitation with 410 nm emission detection. Quantification of SHG collagen signal intensity is shown at right. Scale bar = 200 µm. **(C)** Sircol quantification of soluble collagen secretion normalized to 500,000 cells under 2D and LOL conditions. **(D)** Heatmap of ECM-associated genes in WT, KO and OE conditions in 2D and LOL conditions, shown as WT *vs*. KO and WT *vs*. OE comparisons. **(E)** Heatmap of cell-adhesion-associated gene expression in WT, KO, and OE cells across 2D and LOL conditions, shown as WT *vs.* KO and WT *vs*. OE comparisons. The merged differential expression dataset used to generate these heatmaps is provided in **data file S2**. Western blots represent 5 independent repeats per protein. SHG collagen signal was quantified from 6 spheroids per condition across two independent experiments, with multiple maximum-intensity-projection images per spheroid and multiple ROIs per image; values were averaged per spheroid, with each spheroid treated as one biological replicate (n = 6). Sircol soluble collagen was measured from 6 independent replicates. Data represent mean ± SEM; one-way ANOVA (GraphPad Prism). *p < 0.05, **p < 0.01, ***p < 0.001, ****p < 0.0001; ns, not significant.

To directly assess fibrillar collagen abundance, we performed second-harmonic generation (SHG) microscopy, a label-free technique specific for assembled fibrillar collagen. SHG imaging revealed a marked loss of pericellular and interstitial collagen fibrils in *DMT1* KO spheroids, whereas WT and *DMT1* OE spheroids displayed robust, radially aligned collagen networks **(Fig. 3B and movies S1 to S3)**. Quantitative SHG signal analysis confirmed significantly reduced fibrillar collagen in KO spheroids and increased deposition in OE spheroids (**Fig. 3B**). We next compared these results with Sircol measurements of soluble collagen. Unlike SHG, which detects organized fibrillar collagen, Sircol quantifies total soluble collagen. In 3D LOL cultures, OE showed the highest soluble collagen output, whereas KO remained lower than OE, indicating that soluble collagen abundance and fibrillar collagen assembly do not fully align (**Fig. 3C**). Consistent with these findings, RNA sequencing analyses showed significant downregulation of ECM-associated gene signatures in *DMT1* KO, most prominently in LOL spheroids (**Fig. 3D**). Cell adhesion gene signatures were similarly downregulated in *DMT1* KO cells (**Fig. 3E**). Together, these findings indicate that *DMT1* loss disrupts collagen production and ECM organization, leading us next to determine how these defects affect cancer cell migration and invasive behavior.

### *DMT1* supports cell motility in 2D assays

Iron availability regulates cytoskeletal dynamics (*36*), protease activity, EMT (*37*), and expression of genes required for cell adhesion (*38*). Therefore, we hypothesized that loss of *DMT1* would impair cell motility and invasion in conventional 2D assays. To distinguish planar motility from matrix-embedded invasive behavior, we first assessed *DMT1* WT, KO and OE cells using wound-healing and Transwell Matrigel invasion assays. The wound-healing assay measures collective 2D migration (*39*), whereas the Matrigel invasion assay evaluates the ability of cells to degrade and traverse ECM components (*40–42*). In wound-healing assays in the presence of cytosine arabinoside (Ara-C) to suppress proliferation, *DMT1* KO cells exhibited significantly slower wound closure than WT and OE cells at 24 and 30 h (**Fig. 4A, B**). Similarly, in Transwell Matrigel invasion assays, *DMT1* KO cells showed significantly reduced invasion through Matrigel-coated membranes, whereas WT and OE cells invaded robustly (**Fig. 4C, D**). These findings indicate that *DMT1* loss suppresses migration and invasion in conventional 2D assays. Because these 2D results contrasted with the disrupted spheroid architecture observed in 3D spheroids, we next asked whether *DMT1* loss differentially affects invasive behavior in matrix-embedded LOL tumor spheroids.

**Fig. 4.**
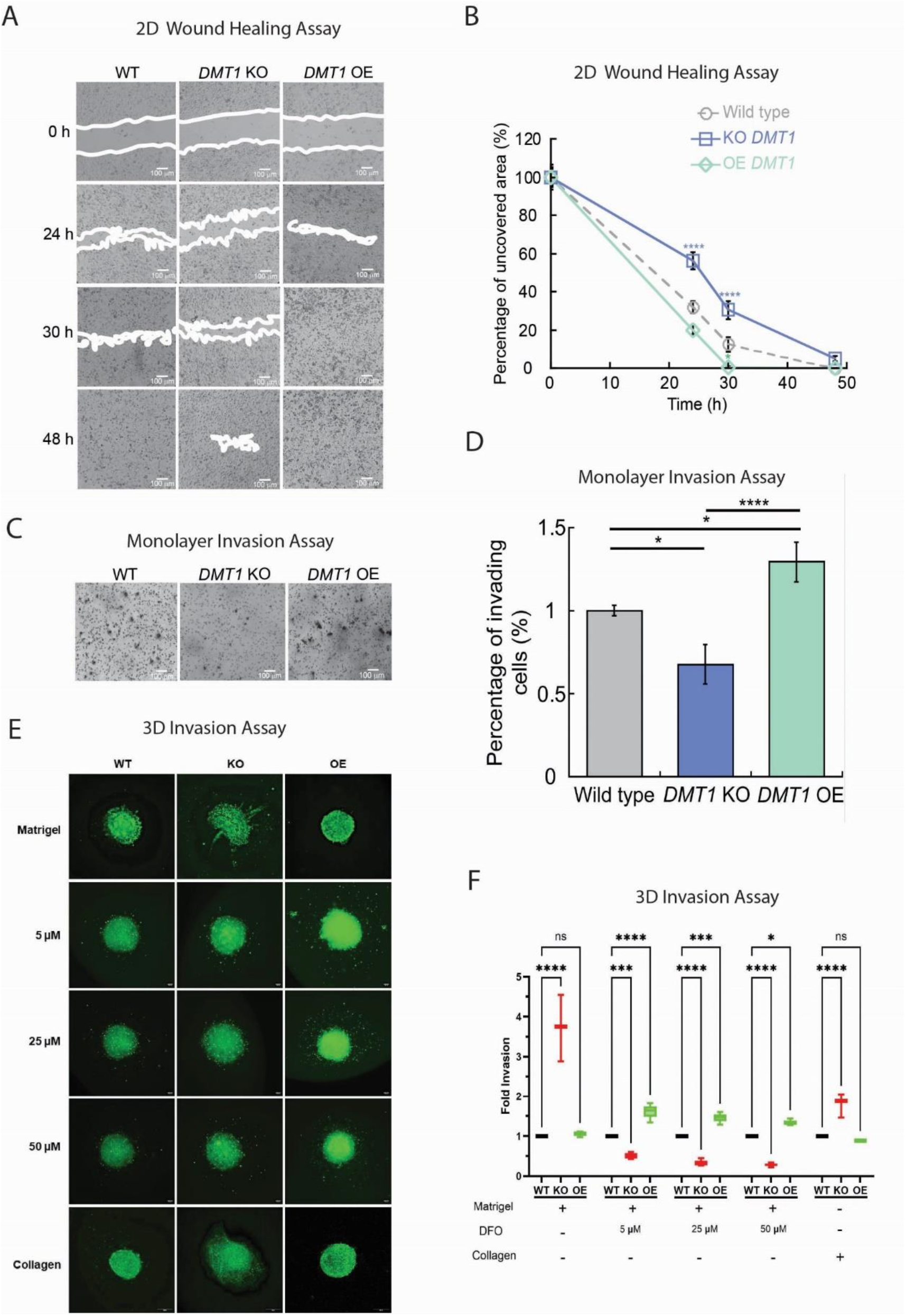
*DMT1* supports motility in 2D but inhibits invasive outgrowth in 3D. **(A, B)** Wound-healing assay showing reduced 2D motility in *DMT1* KO cells relative to WT and *DMT1* OE cells over 48 h. Representative images are shown in **A**, and quantification of uncovered wound area over time is shown in **B**. Scale bar = 100 µm. **(C, D)** Transwell Matrigel invasion assay showing reduced 2D monolayer invasion in *DMT1* KO cells and increased invasion in *DMT1* OE cells relative to WT. Representative images are shown in **C**, and quantification is shown in **D**. Scale bar = 100 µm. **(E, F)** 3D spheroid invasion assay showing invasive outgrowth of WT, *DMT1* KO, and OE LOL spheroids in Matrigel in the absence or presence of deferoxamine (DFO; 5, 25, or 50 µM), and in type I collagen. Representative images are shown in **E**, and quantification of fold invasion is shown in **F**. In the absence of DFO, *DMT1* KO spheroids showed the highest invasive outgrowth. In the presence of DFO, iron chelation suppressed the KO hyperinvasive phenotype, whereas mild iron restriction increased invasive outgrowth in *DMT1* OE spheroids, with limited additional change at higher DFO concentrations. In collagen, *DMT1* KO spheroids again showed the highest invasive capacity. Invasion and migration assays represent 6 independent samples per condition. Data represent mean ± SEM (n = 6); one-way ANOVA (GraphPad Prism). *p < 0.05, **p < 0.01, ***p < 0.001, ****p < 0.0001; ns, not significant.

### *DMT1* loss promotes invasive outgrowth in 3D tumor models

In contrast to the reduced motility observed in conventional 2D assays, we asked whether *DMT1* expression differentially affects invasive behavior in 3D matrices of distinct composition. Because *DMT1* altered both spheroid structural integrity (**Fig. 1**) and ECM organization (**Fig. 3**), we first tested whether these changes translate into altered invasive behavior by embedding intact LOL spheroids in high-density Matrigel (20%) and stimulating them with 40 ng/ml TGF-β1 to induce invasion (*43*). Under these conditions, *DMT1* KO spheroids showed pronounced structural dispersion and invasive outgrowth into the surrounding matrix, whereas *DMT1* OE spheroids retained compact architecture with minimal invasion (**Fig. 4E**). Quantification of fluorescence images showed that, in Matrigel, *DMT1* KO spheroids were significantly more invasive, exhibiting approximately fourfold greater invasive outgrowth into the surrounding matrix than WT and *DMT1* OE spheroids (**Fig. 4F**). These findings reveal a context-dependent phenotype in which *DMT1* loss suppresses motility in 2D yet promotes invasive expansion in 3D.

To determine whether this phenotype depends on matrix composition, we next assessed spheroid invasion in type I collagen. The same pattern was observed in collagen, indicating that the hyperinvasive *DMT1* KO phenotype is not specific to Matrigel. In type I collagen, *DMT1* KO spheroids again showed the highest invasive capacity, whereas *DMT1* OE spheroids remained the least invasive (**Fig. 4E-F**). Across both Matrigel and collagen, KO spheroids consistently lost structural containment, whereas OE spheroids maintained compact spheroid architecture.

Then, we asked whether this 3D invasive phenotype is modulated by iron bioavailability by treating Matrigel-embedded LOL spheroids with increasing concentrations of DFO (5, 25, or 50 µM) added at the time of embedding, with invasion scored from day 4 to day 8 (**Fig. 4E-F**). Iron chelation suppressed the hyperinvasive KO phenotype across the DFO range, whereas mild iron restriction (5 µM DFO) was sufficient to increase invasion in *DMT1* OE spheroids, with minimal additional change at higher DFO concentrations. These genotype-dependent responses indicate that *DMT1* expression state influences how spheroids adapt to altered iron bioavailability and suggest that invasive behavior is governed less by absolute iron abundance than by *DMT1-* dependent iron handling. Together, these findings show that DMT1 loss promotes invasive outgrowth in 3D matrix-embedded spheroid models and that this context-dependent phenotype is further modulated by iron bioavailability. Next, we tested whether ER stress mechanistically connects altered iron distribution to ECM disruption.

### *DMT1* loss induces ER stress and links altered iron homeostasis to ECM disruption

To examine whether the elevated LIP state is associated with ER dysfunction that could contribute to ECM disruption, we analyzed ER-stress pathways. Because accumulation of LIP can perturb the redox and calcium balance required for protein synthesis and folding (*44–46*), we asked whether *DMT1* loss induces ER stress in both 2D and 3D culture models. Western blot analysis and quantification revealed pronounced upregulation of the canonical UPR markers BiP/GRP78 (glucose-regulated protein 78), ATF6 (activating transcription factor 6), and IRE1α (inositol-requiring enzyme 1α) in *DMT1* KO cells grown as 2D monolayers and 3D LOL spheroids (**Fig. 5A**). In contrast, WT and *DMT1* OE cells showed lower levels of these markers, consistent with well-regulated ER homeostasis (**Fig. 5A**).

**Fig. 5.**
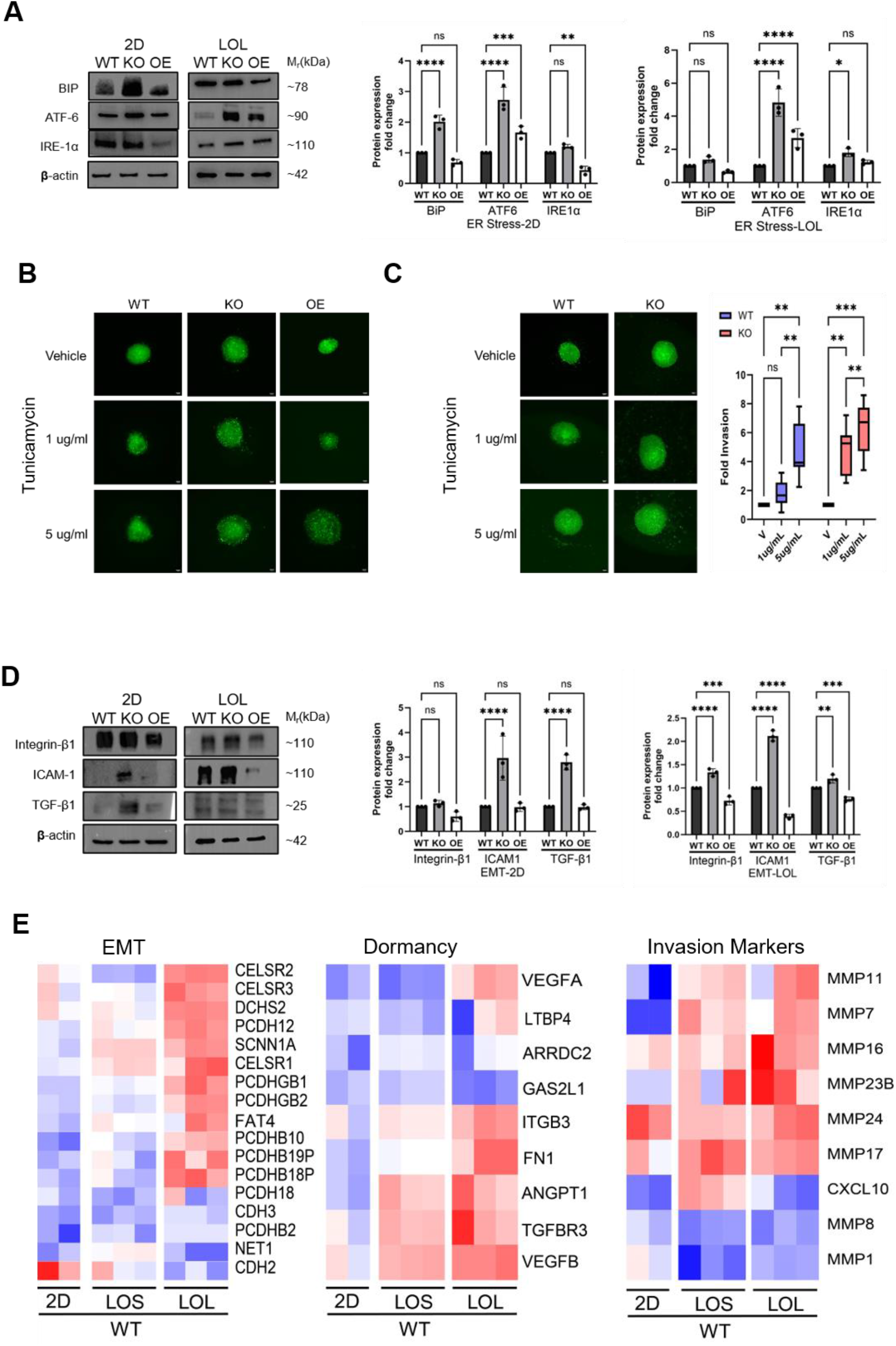
*DMT1* loss is associated with ER stress and EMT-related changes in 3D spheroids. **(A)** Western blot analysis of ER stress/UPR markers (BiP/GRP78, ATF6, and IRE1α) in WT, *DMT1* KO and OE MDA-MB-231 cells grown under 2D and LOL conditions, with β-actin as loading control. Quantification of protein expression in 2D (left) and LOL (right) cultures is shown adjacent to the blots. **(B)** Representative LOL spheroids from WT, KO, and OE cells treated with vehicle, 1 µg/mL tunicamycin, or 5 µg/mL tunicamycin, showing altered spheroid morphology under ER stress-inducing conditions. **(C)** Representative invasion images of tunicamycin-treated spheroids and quantification of fold invasion in WT and *DMT1* KO spheroids under vehicle, 1 µg/mL tunicamycin, and 5 µg/mL tunicamycin conditions. Quantification (right) shows increased invasive area with increasing ER stress. **(D)** Western blot analysis of EMT-related proteins (integrin β1, ICAM1, and TGF-β1) in WT, *DMT1* KO and OE cells under 2D and LOL conditions, with β-actin as loading control. Quantification of protein expression in 2D (left) and LOL (right) cultures is shown adjacent to the blots. **(E)** Heatmaps of representative gene signatures associated with EMT, dormancy, and metastatic hallmarks in WT cells grown in 2D, LOS, and LOL conditions. Western blots represent 5 independent biological replicates per protein; invasion assays represent 6 independent samples per spheroid type. The full list of differentially expressed genes underlying these heatmaps is provided in **data file S1**. Data represent mean ± SEM; one-way ANOVA (GraphPad Prism). *p < 0.05, **p < 0.01, ***p < 0.001, ****p < 0.0001; ns, not significant.

These data suggested that *DMT1* depletion compromises ECM integrity through ER stress activation, prompting us to test whether ER stress is sufficient to disrupt spheroid structure. To this end, WT, *DMT1* KO and OE MDA-MB-231 spheroids were treated with 1 µg/ml or 5 µg/ml tunicamycin, a pharmacological inducer of ER stress and then assessed for functional and invasive phenotypes, as described previously in **Fig. 4E-F** (*47, 48*). Tunicamycin treatment reproduced the *DMT1* KO structural phenotype, as both WT and OE spheroids became loose, irregular, and poorly compacted, resembling untreated *DMT1* KO spheroids (**Fig. 5B**). At higher tunicamycin concentrations (5 µg/ml), WT and KO spheroids showed comparable invasive capacity into the surrounding matrix, indicating that ER stress alone can override DMT1*-*dependent differences in 3D invasion assays (**Fig. 5C**). Together, these findings place ER stress downstream of the DMT1-loss-associated iron imbalance and show that pharmacologic ER stress is sufficient to phenocopy the loss of spheroid compaction and increased 3D invasive outgrowth. Then, we asked whether this ER stress-associated state is accompanied by changes in EMT-related markers that could further explain the enhanced invasive behavior of *DMT1*-depleted spheroids.

### *DMT1* loss-associated ER stress is accompanied by EMT-related changes in 3D tumor spheroids

Given that *DMT1* loss induced ER stress (**Fig. 5A**), and that chronic ER stress can activate EMT through pathways involving TGF-β signaling, PERK-mediated ATF4 activation, and IRE1α-dependent NF-κB signaling (54, 55), we asked whether *DMT1* KO cells exhibit increased EMT-related markers. Therefore, we analyzed the expression of integrin β1, ICAM1 (intercellular adhesion molecule 1), and TGF-β1. In 2D cultures, Western blot analyses of *DMT1* KO cells showed small, non-significant increases in the expression of these three proteins (**Fig. 5D**), while OE cells remained similar to WT.

In 3D culture, however, the pattern was markedly different. *DMT1* KO LOL spheroids showed a clear and significant increase in the EMT related-proteins, integrin β1, ICAM1, and TGF-β1, whereas *DMT1* OE spheroids remained at or slightly below WT levels **(Fig. 5D)**. This dimensionality-dependent effect paralleled the ER stress profile (**Fig. 5A**) and is consistent with a model in which *DMT1* loss promotes an ER stress-associated invasive state under 3D metabolic and hypoxic pressure. To place these protein-level changes in a broader transcriptional context, we examined gene-expression signatures associated with EMT, dormancy, and invasion behavior in WT cells across 2D, LOS, and LOL conditions (**Fig. 5E**). EMT- and metastasis-associated signatures were enriched in LOL spheroids relative to LOS and 2D, whereas dormancy-associated signatures were highest in 2D and progressively decreased in LOS and LOL. These findings indicate that 3D spheroids, particularly LOL, provide a transcriptional context that is more permissive for EMT- and metastasis-associated programs than 2D culture. Because these results linked DMT1-dependent iron and ER stress phenotypes to ECM and EMT-associated changes in 3D tumor models, we next asked whether related relationships could also be detected in patient tumor datasets.

### Patient tumor datasets support coordinated iron-regulatory, ER-proteostasis, and ECM-associated programs

To determine whether components of the experimentally defined iron–ER stress–ECM pathway co-vary in human breast tumors, we performed an exploratory analysis of bulk transcriptomic and proteomic datasets using a 10-marker panel selected a priori to represent the biological modules examined experimentally. Because these datasets contain variable mixtures of malignant, stromal, endothelial, and immune cells, the analysis was not designed to test the proposed tumor-cell mechanism or assign associations to a specific cell population. This panel includes markers of iron regulation (SLC11A2, FTH1), ER stress/proteostasis (HSPA5, P4HB, ERO1A, P4HA1), and ECM/adhesion-associated programs (ITGB1, ICAM1, TGFB1), with HIF1A included based on the hypoxia-related transcriptional context associated with *DMT1* loss in 3D models reported previously by Barra *et al*. (*17*). Details of marker selection, dataset processing, and correlation analysis are given in Materials and Methods.

Because MDA-MB-231 cells model basal-like/TNBC biology, we first examined basal-like tumors in the METABRIC transcriptomic cohort. Spearman correlation analysis revealed a mixed network of positive and negative associations among the 10 markers (ρ range approximately -0.24 to 0.86, complete-case n=198; **Fig. 6A**). Notably, *SLC11A2* (*DMT1*) showed inverse correlations with several stress- and iron-related markers, including HIF1A (ρ = −0.23, p < 0.01), HSPA5 (ρ = −0.22, p < 0.01), ERO1A (ρ = −0.24, p < 0.01), P4HB (ρ = −0.19, p < 0.01), P4HA1 (ρ = −0.17, p < 0.05), and TGFB1 (ρ = −0.15, p < 0.05). In the same matrix, strong positive associations were observed among ER stress/proteostasis markers, including HSPA5–P4HB (ρ = 0.86, p < 0.001), P4HB–P4HA1 (ρ = 0.80, p < 0.001), and HSPA5–ERO1A (ρ = 0.73, p < 0.001), as well as among invasion-related markers such as TGFB1–HIF1A (ρ = 0.56, p < 0.001) and ICAM1–TGFB1 (ρ = 0.54, p < 0.001). These transcriptomic relationships are directionally consistent with coordinated stress- and ECM-associated states in basal-like tumors. These directional relationships are consistent with our experimental model, in which *DMT1* loss is associated with altered iron handling, ER stress, and ECM disruption.

**Fig. 6.**
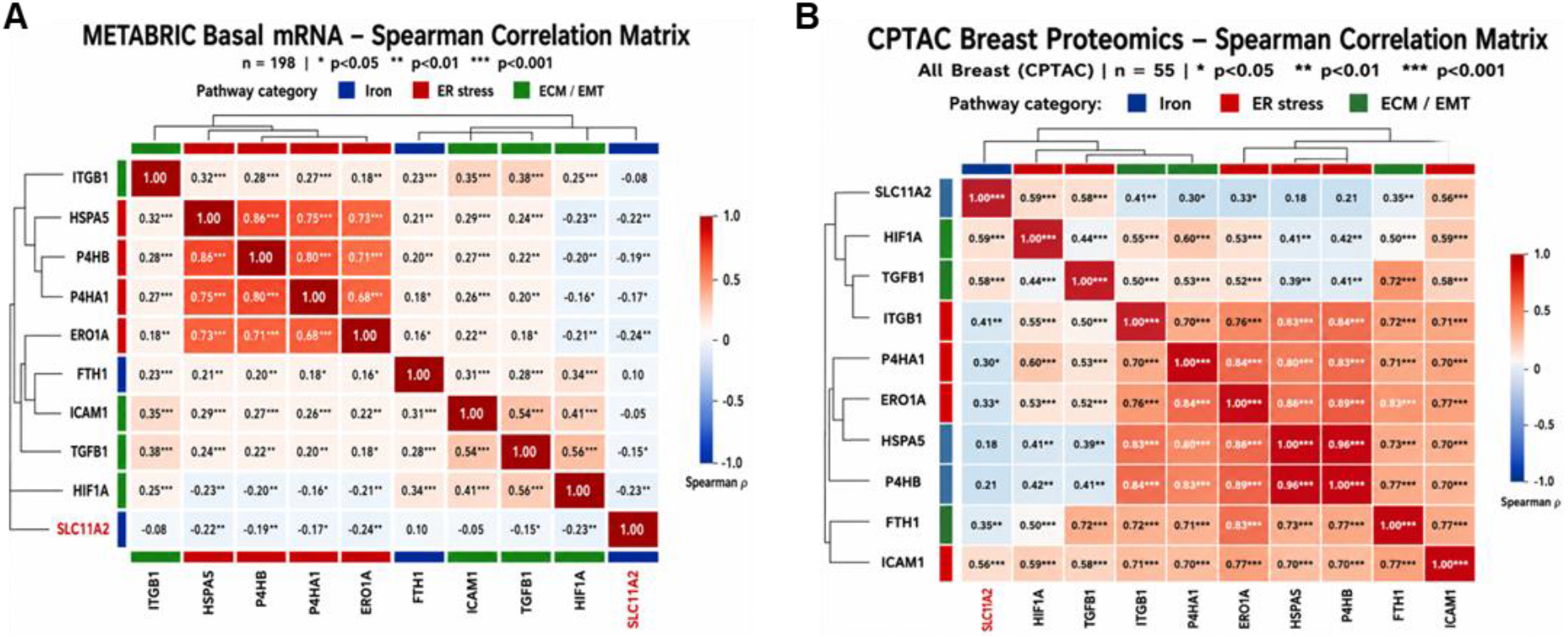
Patient-derived correlation matrices linking iron regulation, ER stress, and ECM/EMT-related programs. **(A)** Spearman correlation matrix of a 10-marker panel in METABRIC transcriptomic dataset from basal-like breast tumors (*n* = 198). The 10-marker panel included SLC11A2 and FTH1 (iron regulation), HSPA5, P4HB, ERO1A, and P4HA1 (ER stress/proteostasis), and ITGB1, ICAM1, TGFB1, and HIF1A (ECM/EMT or hypoxia-associated programs). **(B)** Spearman correlation matrix of the same 10-marker panel in CPTAC breast proteomic samples (*n* = 55). The panel was selected to reflect the biological modules examined in this study, including iron regulation, ER stress/proteostasis, and ECM/EMT or hypoxia-associated programs. Hierarchical clustering was applied to both matrices, and pathway side bars indicate marker category. Cells display Spearman correlation coefficients (ρ). Significance stars denote nominal two-sided p-values (*p<0.05, **p<0.01, ***p<0.001).

We next examined the CPTAC breast proteomics cohort. In contrast to the transcriptomics dataset, protein level Spearman correlations were broadly stronger and predominantly positive (ρ range approximately 0.18 to 0.96, complete-case n=55; **Fig. 6B**). The strongest association in the panel was HSPA5-P4HB (ρ=0.96, p<0.001), followed by strong positive correlations among ER proteostasis and ECM-related proteins, including ERO1A-P4HB (ρ = 0.89, p<0.001), ERO1A-HSPA5 (ρ = 0.86, p<0.001), ITGB1-P4HB (ρ = 0.84, p<0.001), and P4HA1-ERO1A (ρ = 0.84, p<0.001); these results are consistent with coordinated abundance of canonical ER proteostasis components. At the protein level, *SLC11A2* (*DMT1*) showed positive correlations with HIF1A (ρ = 0.59, p<0.001), TGFB1 (ρ = 0.58, p<0.001), ICAM1 (ρ = 0.56, p<0.001), ITGB1 (ρ = 0.41** p<0.001), and FTH1 (ρ = 0.35** p<0.001), whereas its association with HSPA5 was weaker and not significant (ρ = 0.18, ns). These proteomic relationships indicate coordinated abundance across iron-regulatory, ER stress, and ECM-associated modules at the protein level.

Comparison of the 45 shared marker pairs across METABRIC and CPTAC showed substantial overall agreement (Pearson r = 0.67; Spearman ρ = 0.60, computed from the displayed correlation values). Twelve pairs differed in sign between the transcript and protein datasets, and eleven of these involved either SLC11A2 or HIF1A, including SLC11A2–HIF1A (METABRIC ρ = −0.23, p<0.01; CPTAC ρ = 0.59, p<0.001) and ERO1A–HIF1A (METABRIC ρ = −0.21, p<0.01; CPTAC ρ = 0.53, p<0.001). Associations among the ER proteostasis and ECM markers were positive and of similar rank in both cohorts.

## DISCUSSION

DMT1 delivers transferrin-derived iron from endosomes to mitochondria by supporting endosome–mitochondria contacts, and its loss redirects that iron into an expanded labile pool while increasing metastatic outgrowth (*17*). We propose that this redistribution is not simply a change in where iron sits, but a change in which compartments can use it. In DMT1 KO cells, total iron falls while the labile pool roughly doubles, a combination that rules out uniform iron deficiency and points instead to defective intracellular partitioning. This pattern is not unique to our system. Misdistribution of iron between mitochondria and cytosol, rather than absolute deficiency, underlies the cellular defects of frataxin deficiency, and relocating iron between compartments restores function without altering total iron content (*49*). In PKAN astrocytes, fewer transferrin-containing vesicles contact mitochondria, mitochondrial iron availability falls, and the cell becomes cytosolically iron-loaded, the same directional split we observe after DMT1 loss (*50*). The functional consequence runs in two directions at once: redox-active Fe²⁺ accumulates in compartments where it drives stress, while iron-dependent biosynthetic reactions in the secretory pathway lose access to the substrate they require. That second arm bears directly on ECM output, because the collagen prolyl 4-hydroxylases and lysyl hydroxylases are ER-luminal Fe²⁺-dependent dioxygenases whose activity depends on iron delivered to the secretory pathway rather than on bulk cellular iron (*51*). ER stress and impaired collagen output are the predicted readouts of that split, and both are what we observe. Within this model, iron availability to specific organelles, rather than bulk iron abundance, is what determines ECM integrity and invasive behavior.

The increase in LIP in *DMT1*-deficient cells was accompanied by broad activation of the UPR pathway, with increased BiP/GRP78, ATF6, and IRE1α in both 2D and 3D models. These findings support the idea that compartmentalized iron imbalance compromises ER proteostasis, rather than activating a selective stress branch (*45, 52*). Reduced ERO1 further suggests impaired oxidative folding capacity, consistent with defective maturation of secreted proteins such as collagen (*53, 54*). In this context, expansion of the labile iron pool does not necessarily indicate adequate iron availability to ER-associated biosynthetic pathways; instead, redox-active iron may accumulate in inappropriate intracellular pools while iron-dependent collagen-processing functions remain limited. The collagen phenotype was supported by multiple readouts: *DMT1* loss reduced COL1A1, disrupted fibrillar collagen organization by SHG, altered soluble collagen output, and downregulated ECM- and adhesion-related transcriptional programs, supporting the concept that iron dysregulation translates into ECM disorganization. Together, these data support a model in which altered intracellular iron distribution compromises collagen production and ECM organization through ER stress-associated defects in biosynthetic capacity.

Pharmacologic ER stress induction independently reproduced the loose spheroid architecture and invasive phenotype, indicating that ER dysfunction is not merely associated with, but functionally sufficient to phenocopy key features of ECM destabilization. These findings support a model in which ER stress acts downstream of DMT1*-*dependent iron imbalance and is sufficient to reproduce major aspects of the ECM and invasion phenotype in TNBC (*48, 55*). Chelation experiments revealed that *DMT1* genotype shapes invasion responses to altered iron availability, suggesting that invasive behavior depends on intracellular iron distribution rather than on iron abundance alone. In particular, DFO suppressed the hyperinvasive phenotype of *DMT1* KO spheroids but increased invasion in DMT1-overexpressing spheroids at low concentrations, indicating that the response to iron restriction is determined by DMT1-dependent iron handling rather than by a uniform relationship between iron depletion and invasion. The convergence of genetic and pharmacologic disruption of iron homeostasis on altered invasive behavior further supports intracellular iron partitioning, rather than simple deficiency, as a regulator of ECM integrity and cancer cell invasion.

Beyond proteostasis and ECM destabilization, *DMT1* deficiency appears to shift cancer cells from an ECM-maintaining state toward a stress-adapted invasive program. Importantly, the consequences of *DMT1* loss were strongly context dependent: in conventional 2D assays, *DMT1* loss reduced wound closure and Transwell Matrigel invasion, whereas in 3D matrix-embedded spheroids it promoted loose aggregate formation, loss of structural containment, and increased invasive outgrowth in both Matrigel and collagen matrices. The divergence between reduced 2D motility and enhanced 3D invasive outgrowth indicates that *DMT1* loss primarily alters tumor structural adaptation and ECM remodeling rather than intrinsic planar migration alone (*56*). Moreover, this context dependence is consistent with the idea that altered iron distribution influences both structural matrix integrity and broader transcriptional plasticity, which is consistent with a role for iron-responsive elements (IREs) and iron regulatory proteins (IRPs) in post-transcriptional gene regulation (*57, 58*), although the precise regulatory mechanisms linking these states remain to be defined (*59*). Bulk RNA sequencing analyses further showed that *DMT1* loss decreased the gene expression of genes related to ECM maintenance and cell adhesion including *COL1A1, COL3A1*, fibronectin, and laminins. Consistently, *DMT1* KO cells showed upregulation of EMT-associated genes, accompanied by increased expression of integrin β1, ICAM1, and TGF-β1 at the protein level (*60*). The strong induction of TGF-β1 highlights its central role in driving EMT transcriptional programs, while increased integrin β1 and ICAM1 are consistent with a stress-associated invasive state in 3D environments. Together with ECM disruption, these changes support a model where *DMT1* KO cells show enhanced invasive properties in both the collagen and Matrigel 3D matrices. Conversely, *DMT1* OE maintained spheroid compactness and reduced invasion, consistent with a protective role for *DMT1* in preserving structural stability under 3D conditions. In summary, *DMT1* loss and subsequent disrupted iron distribution does not uniformly enhance motility; rather, it selectively promotes invasive behavior in 3D environments where ECM organization, multicellular architecture, and structural confinement become dominant constraints.

To assess whether the relationships identified in our model extend to human disease, we analyzed patient tumor datasets for coordinated iron-, ER stress-, and ECM-related programs. The METABRIC basal-like transcriptomic cohort provided the closer directional match to our experimental model: lower *SLC11A2* expression correlated with higher levels of *HIF1A*, *HSPA5*, *ERO1A*, *P4HB*, *P4HA1*, and *TGFB1*, consistent with coordinated stress- and ECM-associated transcriptional states in tumors with lower *DMT1* expression. By contrast, the CPTAC proteomic dataset showed broadly stronger and predominantly positive co-abundance relationships across the same marker panel. This difference does not necessarily contradict the experimental findings, but it changes how the proteomic analysis should be interpreted. Our western blot analyses reflect genotype-driven changes in a controlled perturbation system, whereas CPTAC reflects bulk-tumor protein co-abundance across heterogeneous human samples. Accordingly, the CPTAC analysis is best viewed as evidence that iron-regulatory, ER stress, and ECM-related programs remain coupled at the protein level in human tumors, rather than as a direct directional validation of the *DMT1*-loss phenotype. The correlation structures of the two cohorts agree closely overall, with the discordance confined almost entirely to SLC11A2 and HIF1A. Two differences between the analyses likely contribute: the METABRIC panel is restricted to basal-like tumors whereas the CPTAC panel spans all breast subtypes, and bulk protein co-abundance is shaped by cell-type composition and protein stability in ways that transcript abundance is not.

Our conclusions rest on an isogenic MDA-MB-231 series, and the compartmentalization model is inferred rather than measured, since FerroOrange-based labile iron reports chelatable cytosolic iron and does not resolve the pool available to ER-luminal hydroxylases. Tunicamycin reproduced the loose spheroid architecture and invasive outgrowth, which establishes that ER stress is sufficient for the phenotype but not that DMT1 acts exclusively through it, and the chelation data derive from a single agent over a limited concentration range. The patient associations are drawn from bulk cohorts and show that these programs co-vary in human tumors without assigning causality or reporting subcellular iron. Extending this to patients would begin with assessment of ER stress and collagen organization in triple-negative tumors stratified by DMT1 expression, followed by in vivo work asking whether clinically available chelators suppress or enhance metastatic outgrowth as a function of DMT1 status. A marker scoreable in archival tissue, with a threshold set against outcome, would be needed before stratification by tumor iron handling could be tested prospectively.

Overall, our findings define an iron-ER-ECM axis, schematically illustrated in **Fig. 7**, in which *DMT1-*dependent intracellular iron partitioning supports ER proteostasis, collagen production, and ECM integrity, thereby restraining invasive outgrowth in 3D tumor models. The central implication is that total cellular iron does not predict invasive behavior without considering its intracellular distribution and functional availability to specific organelles and biosynthetic pathways. Our data further identify ER stress as a plausible mechanistic bridge between altered iron homeostasis and collagen synthesis failure. These findings may help explain why lowering bulk iron does not necessarily suppress invasive phenotypes uniformly and instead suggest that therapeutic responses may depend on how iron trafficking pathways reshape ER proteostasis, ECM organization, and tumor architecture.

**Fig. 7.**
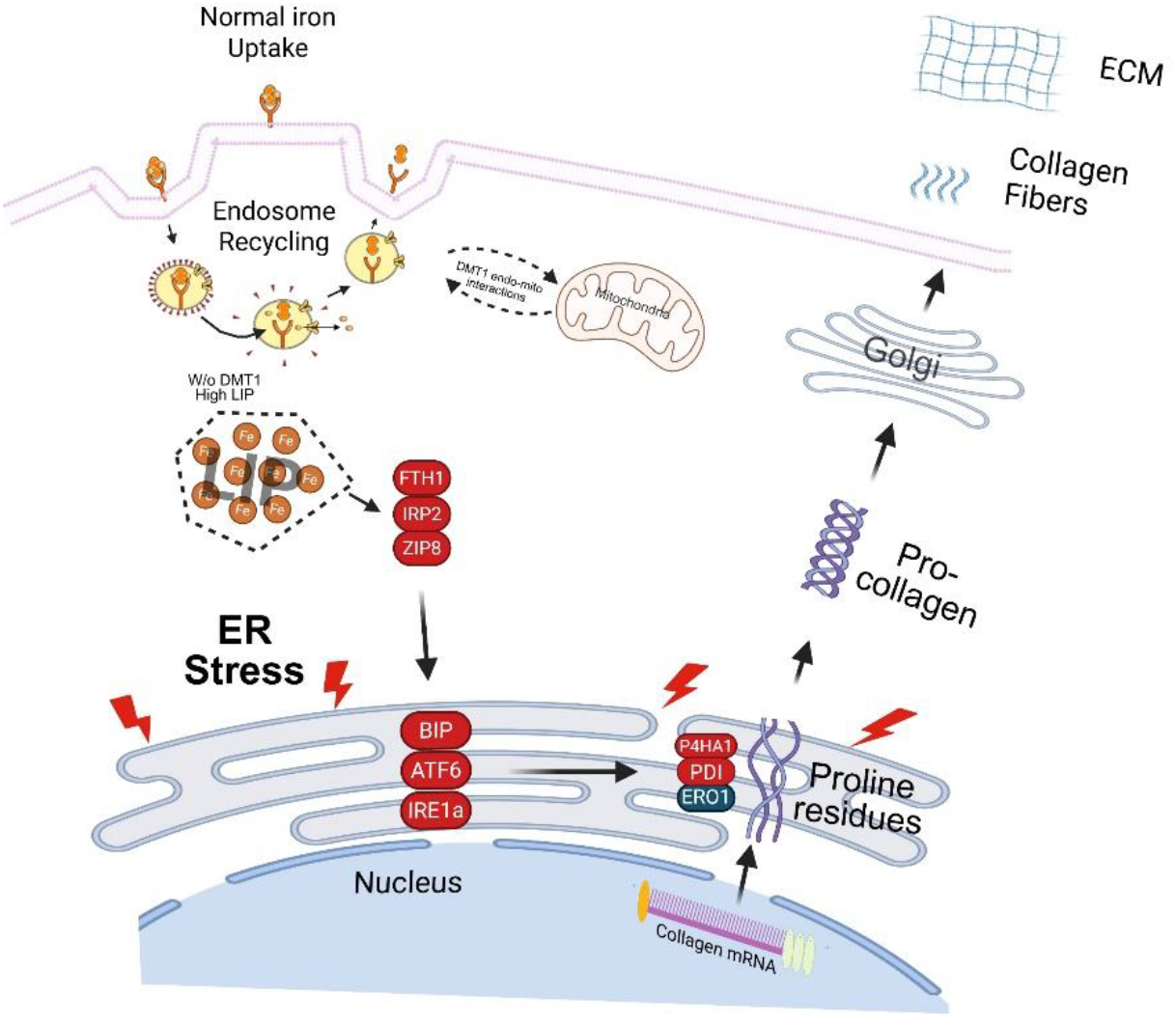
Schematic model of the *DMT1*-iron-ER stress-ECM axis. Proposed model summarizing how *DMT1* loss alters intracellular iron homeostasis in TNBC cells. Although total iron content is reduced, *DMT1* loss is associated with expansion of the labile iron pool (LIP), activation of ER stress/UPR markers (BiP/GRP78, ATF6, IRE1α), and disruption of collagen biosynthesis and maturation pathways. Reduced ERO1 together with altered PDI/P4HB and P4HA1-associated processing is proposed to impair procollagen maturation, resulting in defective collagen organization and disrupted ECM architecture. These changes are associated with loss of spheroid structural containment and increased cancer cell invasion in 3D tumor models. Red ovals, upregulated proteins; blue ovals, downregulated proteins; lightning bolts, ER stress; orange circles, Fe²⁺. Figure created with Biorender (agreement # IW29RHRJ5H).

## MATERIALS AND METHODS

### Study design

This study tested whether DMT1-dependent intracellular iron distribution regulates extracellular matrix organization and invasive behavior in triple-negative breast cancer, and whether those effects depend on culture dimensionality. We used a previously generated isogenic MDA-MB-231 panel comprising wild-type (WT), *DMT1* knockout (KO), and *DMT1*-GFP overexpressing (OE) cells (*17*), compared in parallel as 2D monolayers and as liquid-overlay 3D spheroids at two defined sizes. Prospectively defined endpoints were total cellular iron, labile Fe²⁺ pool, collagen abundance and fibrillar organization, unfolded protein response marker levels, and invasive outgrowth into Matrigel and type I collagen. Iron availability was manipulated with deferoxamine and ER stress was induced independently with tunicamycin to test sufficiency and directionality. Imaging endpoints were quantified from six spheroids per condition across two independent experiments, with each spheroid treated as one biological replicate; immunoblots represent five independent biological replicates per protein, and invasion assays six independent samples per condition. Replicate numbers for individual panels are stated in the figure legends. Experiments were not randomized, and investigators were not blinded during acquisition or analysis, as cell genotypes were identified by constitutive fluorescent reporters. No data were excluded. Relationships identified in the isogenic panel were then tested for concordance in human breast tumor transcriptomic (METABRIC) and proteomic (CPTAC) cohorts.

### Cell culture

The MDA-MB-231 cell line was obtained from ATCC (HTB-26), and the DMT1 knockout (KO) and DMT1-OE-GFP overexpression (OE) variants were generated in-house as described previously (*17*). All lines were maintained under identical conditions at 37 °C and 5% CO₂ in humidified incubators using standard or phenol red–free DMEM supplemented with 10% fetal bovine serum, 100 U/ml penicillin, and 100 µg/ml streptomycin. Functional uptake, endosomal trafficking, and intracellular iron release of human iron-loaded transferrin in MDA-MB-231 cells, as well as transferrin uptake throughout MDA-MB-231 spheroids, have been demonstrated previously (*14, 16, 17*). Cultures were screened regularly for mycoplasma by PCR and used at passage 10 or lower. Reagents, antibodies, cell lines, and software used throughout the study are listed in **table S1**.

### Liquid overlay spheroid formation and analysis

Multicellular tumor spheroids were generated by the liquid overlay method in non-adherent 96-well U-bottom plates (CellStar, Greiner Bio-One). Monolayers were trypsinized, counted on a CellDrop automated counter (DeNovix), and resuspended in complete DMEM. Two spheroid sizes were produced by varying the cell seeding density, 25,000 cells per well for large (LOL) and 8,000 cells per well for small (LOS) spheroids. LOS spheroids measured approximately 300 to 500 µm in diameter and LOL spheroids approximately 500 to 800 µm. Each well received 50 µl of cell suspension layered over 10% Matrigel® in DMEM, giving a final concentration of 5% Matrigel® (*61*). Plates were centrifuged at 200 × *g* for 10 min to promote aggregation and then cultured for 4 days. Unless noted otherwise, all experiments used 4-day spheroids, except the 3D invasion assay. Spheroid morphology was assessed on day 4 by phase-contrast imaging on a Leica Thunder DMi8 microscope (Leica Microsystems) with an N PLAN 5×/0.12 PH0 dry objective.

### Data and statistical analysis

Ungrouped datasets were analyzed by one-way ANOVA and grouped datasets by two-way ANOVA. Data are shown as mean ± SEM. All experiments were performed at least in triplicate, with cell numbers for imaging analyses given in the figure legends. P < 0.05 was considered significant (*p < 0.05; **p < 0.01). Analysis and plotting were done in GraphPad Prism 10.0 (Dotmatics).

Detailed procedures for all other assays are described in Materials and Methods in the Supplementary Materials.

### List of Supplementary Materials

## Materials and Methods

Fig. S1 to S5

Tables S1 to S2

Data files S1 to S3

Movies S1 to S3

References (62–85)

## Acknowledgments

We would like to acknowledge the support of the AMC imaging core for the use of the LSM 880 confocal microscope, Thunder microscope and Imaris software. We would like to thank the NIH and the 1S10OD034442 shared instrumentation award for the use of the Nikon AXR-MP multiphoton microscope.

## Author contributions

Conceptualization: AA, MB

Methodology: AA, PS, RBR, DTC, TPB, MB

Investigation: AA, KP, KN, PS, OD, IC, JD, TH, RBR, TPB

Formal analysis: AA, RBR, DTC, TPB, MB Resources: DTC, TPB, MB

Data curation: AA,

Visualization: AA, PS, DTC, MB

Supervision: DTC, TPB, MB

Project administration: MB, DTC, TPB

Funding acquisition: MB, DTC, TPB

Writing – original draft: AA, TPB, MB

Writing – review & editing: AA, KP, KN, PS, OD, IC, JD, TH, RBR, DTC, TPB, MB

## Data and materials availability

RNA sequencing data generated in this study (MDA-MB-231 WT and *DMT1* OE cells across 2D, LOS, and LOL conditions) have been deposited in the NCBI Gene Expression Omnibus under accession GSE312425. Previously published RNA sequencing data for MDA-MB-231 WT and *DMT1* KO cells (*17*) are available under accession GSE226059. Patient transcriptomic data were obtained from the METABRIC cohort via cBioPortal, and proteomic data from the CPTAC breast cancer cohort via the NCI Proteomic Data Commons.

## References and Notes

1. Y. Chen, Z. Fan, Y. Yang, C. Gu, Iron metabolism and its contribution to cancer. International journal of oncology 54, 1143–1154 (2019).

2. W. P. Faulk, B. L. Hsi, P. J. Stevens, Transferrin and transferrin receptors in carcinoma of the breast.Lancet 2, 390–392 (1980).

3. S. V. Torti, F. M. Torti, Iron and cancer: more ore to be mined. Nature Reviews Cancer 13, 342–355 (2013).

4. S. J. Dixon, B. R. Stockwell, The role of iron and reactive oxygen species in cell death. Nature Chemical Biology 10, 9–17 (2014).

5. M. Huang, Y. Wang, X. Wu, W. Li, Crosstalk between Endoplasmic Reticulum Stress and Ferroptosis in Liver Diseases. FBL 29, (2024).

6. T. C. H. Tan, D. H. G. Crawford, L. A. Jaskowski, V. N. Subramaniam, A. D. Clouston, D. I. Crane, K. R. Bridle, G. J. Anderson, L. M. Fletcher, Excess iron modulates endoplasmic reticulum stress-associated pathways in a mouse model of alcohol and high-fat diet-induced liver injury. Laboratory Investigation 93, 1295–1312 (2013).

7. B. Galy, M. Conrad, M. Muckenthaler, Mechanisms controlling cellular and systemic iron homeostasis.Nature Reviews Molecular Cell Biology 25, 133–155 (2024).

8. A. Asif, K. H. Kim, F. Jabbar, S. Kim, K. H. Choi, Real-time sensors for live monitoring of disease and drug analysis in microfluidic model of proximal tubule. Microfluidics and Nanofluidics 24, 43 (2020).

9. A. Asif, S. H. Park, A. Manzoor Soomro, M. A. U. Khalid, A. R. C. Salih, B. Kang, F. Ahmed, K. H. Kim, K. H. Choi, Microphysiological system with continuous analysis of albumin for hepatotoxicity modeling and drug screening. Journal of Industrial and Engineering Chemistry 98, 318–326 (2021).

10. M. Domingues, C. Leite Pereira, B. Sarmento, F. Castro, Mimicking 3D breast tumor-stromal interactions to screen novel cancer therapeutics. European Journal of Pharmaceutical Sciences 190, 106560 (2023).

11. A. S. Nunes, A. S. Barros, E. C. Costa, A. F. Moreira, I. J. Correia, 3D tumor spheroids as in vitro models to mimic in vivo human solid tumors resistance to therapeutic drugs. Biotechnology and Bioengineering 116, 206–226 (2019).

12. N. Hubert, M. W. Hentze, Previously uncharacterized isoforms of divalent metal transporter (DMT)-1: Implications for regulation and cellular function. Proceedings of the National Academy of Sciences 99, 12345–12350 (2002).

13. H. Gunshin, B. Mackenzie, U. V. Berger, Y. Gunshin, M. F. Romero, W. F. Boron, S. Nussberger, J. L. Gollan, M. A. Hediger, Cloning and characterization of a mammalian proton-coupled metal-ion transporter. Nature 388, 482–488 (1997).

14. T. C. Khoo, K. Tubbesing, A. Rudkouskaya, S. Rajoria, A. Sharikova, M. Barroso, A. Khmaladze, Quantitative label-free imaging of iron-bound transferrin in breast cancer cells and tumors. Redox Biol 36, 101617 (2020).

15. K. Tubbesing, J. Ward, R. Abini-Agbomson, A. Malhotra, A. Rudkouskaya, J. Warren, J. Lamar, N. Martino, A. P. Adam, M. Barroso, Complex Rab4-Mediated Regulation of Endosomal Size and EGFR Activation. Mol Cancer Res 18, 757–773 (2020).

16. D. M. Kingsley, C. L. Roberge, A. Rudkouskaya, D. E. Faulkner, M. Barroso, X. Intes, D. T. Corr, Laser-based 3D bioprinting for spatial and size control of tumor spheroids and embryoid bodies. Acta Biomater 95, 357–370 (2019).

17. J. Barra, I. Crosbourne, C. L. Roberge, R. Bossardi-Ramos, J. S. A. Warren, K. Matteson, L. Wang, F. Jourd’heuil, S. M. Borisov, E. Bresnahan, J. J. Bravo-Cordero, R. I. Dmitriev, D. Jourd’heuil, A. P. Adam, J. M. Lamar, D. T. Corr, M. M. Barroso, DMT1-dependent endosome-mitochondria interactions regulate mitochondrial iron translocation and metastatic outgrowth. Oncogene 43, 650–667 (2024).

18. N. A. Wolff, M. D. Garrick, L. Zhao, L. M. Garrick, A. J. Ghio, F. Thévenod, A role for divalent metal transporter (DMT1) in mitochondrial uptake of iron and manganese. Sci Rep 8, 211 (2018).

19. X. Xue, Sadeesh K. Ramakrishnan, K. Weisz, D. Triner, L. Xie, D. Attili, A. Pant, B. Győrffy, M. Zhan, C. Carter-Su, Karin M. Hardiman, Thomas D. Wang, Michael K. Dame, J. Varani, D. Brenner, Eric R. Fearon, Yatrik M. Shah, Iron Uptake via DMT1 Integrates Cell Cycle with JAK-STAT3 Signaling to Promote Colorectal Tumorigenesis. Cell Metabolism 24, 447–461 (2016).

20. T. Hoki, E. Katsuta, L. Yan, K. Takabe, F. Ito, Low DMT1 Expression Associates With Increased Oxidative Phosphorylation and Early Recurrence in Hepatocellular Carcinoma. Journal of Surgical Research 234, 343–352 (2019).

21. P. Lelièvre, L. Sancey, J.-L. Coll, A. Deniaud, B. Busser, Iron Dysregulation in Human Cancer: Altered Metabolism, Biomarkers for Diagnosis, Prognosis, Monitoring and Rationale for Therapy. Cancers 12, 3524 (2020).

22. H. Zhang, N. Chen, C. Ding, H. Zhang, D. Liu, S. Liu, Ferroptosis and EMT resistance in cancer: a comprehensive review of the interplay. Frontiers in Oncology **Volume** 14 **-** 2024, (2024).

23. F. Kai, A. P. Drain, V. M. Weaver, The extracellular matrix modulates the metastatic journey. Developmental cell 49, 332–346 (2019).

24. R. Borst, L. Meyaard, M. I. Pascoal Ramos, Understanding the matrix: collagen modifications in tumors and their implications for immunotherapy. Journal of Translational Medicine 22, 382 (2024).

25. Y. Wang, L. Yu, J. Ding, Y. Chen, Iron Metabolism in Cancer. International Journal of Molecular Sciences 20, 95 (2019).

26. R. Yu, Y. Hang, H.-i. Tsai, D. Wang, H. Zhu, Iron metabolism: backfire of cancer cell stemness and therapeutic modalities. Cancer Cell International 24, 157 (2024).

27. P. Walter, D. Ron, The Unfolded Protein Response: From Stress Pathway to Homeostatic Regulation. Science 334, 1081–1086 (2011).

28. S. Hisanaga, M. Miyake, S. Taniuchi, M. Oyadomari, M. Morimoto, R. Sato, J. Hirose, H. Mizuta, S. Oyadomari, PERK-mediated translational control is required for collagen secretion in chondrocytes. Sci Rep 8, 773 (2018).

29. S. Zhu, J. Yin, X. Lu, D. Jiang, R. Chen, K. Cui, W. He, N. Huang, G. Xu, Influence of experimental variables on spheroid attributes. Scientific Reports 15, 9751 (2025).

30. M. Rovere, D. Reverberi, P. Arnaldi, M. E. F. Palamà, C. Gentili, Spheroid size influences cellular senescence and angiogenic potential of mesenchymal stromal cell-derived soluble factors and extracellular vesicles. Frontiers in Bioengineering and Biotechnology **Volume** 11 - 2023, (2023).

31. K. Froehlich, J.-D. Haeger, J. Heger, J. Pastuschek, S. M. Photini, Y. Yan, A. Lupp, C. Pfarrer, R. Mrowka, E. Schleußner, U. R. Markert, A. Schmidt, Generation of Multicellular Breast Cancer Tumor Spheroids: Comparison of Different Protocols. Journal of Mammary Gland Biology and Neoplasia 21, 89–98 (2016).

32. C. L. Roberge, D. M. Kingsley, D. E. Faulkner, C. J. Sloat, L. Wang, M. Barroso, X. Intes, D. T. Corr, Non-Destructive Tumor Aggregate Morphology and Viability Quantification at Cellular Resolution, During Development and in Response to Drug. Acta Biomaterialia 117, 322–334 (2020).

33. S. J. Gordon, Y. Xiao, A. L. Paskavitz, N. Navarro-Tito, J. G. Navea, T. Padilla-Benavides, Atomic absorbance spectroscopy to measure intracellular zinc pools in mammalian cells. Journal of Visualized Experiments (JoVE), e59519 (2019).

34. C. McCann, M. Quinteros, I. Adelugba, M. N. Morgada, A. R. Castelblanco, E. J. Davis, A. Lanzirotti, S. J. Hainer, A. J. Vila, J. G. Navea, T. Padilla-Benavides, The mitochondrial Cu+ transporter PiC2 (SLC25A3) is a target of MTF1 and contributes to the development of skeletal muscle in vitro. Frontiers in Molecular Biosciences **Volume** 9 - 2022, (2022).

35. S. J. V. Gordon, D. E. Fenker, K. E. Vest, T. Padilla-Benavides, Manganese influx and expression of ZIP8 is essential in primary myoblasts and contributes to activation of SOD2†. Metallomics 11, 1140–1153 (2019).

36. A. N. Goldfarb, K. C. Freeman, R. K. Sahu, K. E. Elagib, M. Holy, A. Arneja, R. Polanowska-Grabowska, A. A. Gru, Z. White, S. Khalil, M. J. Kerins, A. Ooi, N. Leitinger, C. J. Luckey, L. L. Delehanty, Iron control of erythroid microtubule cytoskeleton as a potential target in treatment of iron-restricted anemia. Nature Communications 12, 1645 (2021).

37. Y. Han, L. Ye, F. Du, M. Ye, C. Li, X. Zhu, Q. Wang, H. Jiang, Z. Liu, J. Ma, J. Zhou, C. Bai, Y. Song, J. Liu, Iron metabolism regulation of epithelial-mesenchymal transition in idiopathic pulmonary fibrosis. Annals of Translational Medicine 9, 1755 (2021).

38. A. E. R. Kartikasari, N. A. Georgiou, F. L. J. Visseren, H. van Kats-Renaud, B. S. van Asbeck, J. J. M. Marx, Intracellular Labile Iron Modulates Adhesion of Human Monocytes to Human Endothelial Cells. *Arteriosclerosis*, Thrombosis, and Vascular Biology 24, 2257–2262 (2004).

39. A. V. P. Bobadilla, J. Arévalo, E. Sarró, H. M. Byrne, P. K. Maini, T. Carraro, S. Balocco, A. Meseguer, T. Alarcón, *In vitro* cell migration quantification method for scratch assays. Journal of The Royal Society Interface 16, 20180709 (2019).

40. E. Y. Kim, O. Verdejo-Torres, K. Diaz-Rodriguez, F. Hasanain, L. Caromile, T. Padilla-Benavides, Single nucleotide polymorphisms and Zn transport by ZIP11 shape functional phenotypes of HeLa cells. Metallomics 16, (2024).

41. M. Olea-Flores, J. Kan, A. Carlson, S. A. Syed, C. McCann, V. Mondal, C. Szady, H. M. Ricker, A. McQueen, J. G. Navea, L. A. Caromile, T. Padilla-Benavides, ZIP11 Regulates Nuclear Zinc Homeostasis in HeLa Cells and Is Required for Proliferation and Establishment of the Carcinogenic Phenotype. Front Cell Dev Biol 10, 895433 (2022).

42. M.-J. Park, H.-J. Kwak, H.-C. Lee, D.-H. Yoo, I.-C. Park, M.-S. Kim, S.-H. Lee, C. H. Rhee, S.-I. Hong, Nerve Growth Factor Induces Endothelial Cell Invasion and Cord Formation by Promoting Matrix Metalloproteinase-2 Expression through the Phosphatidylinositol 3-Kinase/Akt Signaling Pathway and AP-2 Transcription Factor*. Journal of Biological Chemistry 282, 30485–30496 (2007).

43. Y. Drabsch, P. Ten Dijke, TGF-β signaling in breast cancer cell invasion and bone metastasis. Journal of mammary gland biology and neoplasia 16, 97–108 (2011).

44. S. S. Cao, R. J. Kaufman, Endoplasmic reticulum stress and oxidative stress in cell fate decision and human disease. Antioxid Redox Signal 21, 396–413 (2014).

45. J. Che, H. Lv, J. Yang, B. Zhao, S. Zhou, T. Yu, P. Shang, Iron overload induces apoptosis of osteoblast cells via eliciting ER stress-mediated mitochondrial dysfunction and p-eIF2α/ATF4/CHOP pathway in vitro. Cellular Signalling 84, 110024 (2021).

46. T. Makio, J. Chen, T. Simmen, ER stress as a sentinel mechanism for ER Ca2+ homeostasis. Cell Calcium 124, 102961 (2024).

47. Z. You, L. He, N. Yan, Tunicamycin-Induced Endoplasmic Reticulum Stress Promotes Breast Cancer Cell MDA-MB-231 Apoptosis through Inhibiting Wnt/β-Catenin Signaling Pathway. Journal of Healthcare Engineering 2021, 6394514 (2021).

48. J. Wu, S. Chen, H. Liu, Z. Zhang, Z. Ni, J. Chen, Z. Yang, Y. Nie, D. Fan, Tunicamycin specifically aggravates ER stress and overcomes chemoresistance in multidrug-resistant gastric cancer cells by inhibiting N-glycosylation. Journal of Experimental & Clinical Cancer Research 37, 272 (2018).

49. O. Kakhlon, H. Manning, W. Breuer, N. Melamed-Book, C. Lu, G. Cortopassi, A. Munnich, Z. I. Cabantchik, Cell functions impaired by frataxin deficiency are restored by drug-mediated iron relocation. Blood 112, 5219–5227 (2008).

50. P. Santambrogio, A. Cozzi, C. Balestrucci, M. Ripamonti, V. Berno, E. Cammarota, A. S. Moro, S. Levi, Mitochondrial iron deficiency triggers cytosolic iron overload in PKAN hiPS-derived astrocytes. Cell Death Dis 15, 361 (2024).

51. A. M. Salo, J. Myllyharju, Prolyl and lysyl hydroxylases in collagen synthesis. Exp Dermatol 30, 38–49 (2021).

52. M. Halon-Golabek, A. Borkowska, A. Herman-Antosiewicz, J. Antosiewicz, Iron Metabolism of the Skeletal Muscle and Neurodegeneration. Frontiers in Neuroscience **Volume** 13 - 2019, (2019).

53. N. J. Bulleid, L. Ellgaard, Multiple ways to make disulfides. Trends in Biochemical Sciences 36, 485–492 (2011).

54. C. S. Sevier, C. A. Kaiser, Formation and transfer of disulphide bonds in living cells. Nature Reviews Molecular Cell Biology 3, 836–847 (2002).

55. S. You, W. Li, Y. Guan, Tunicamycin inhibits colon carcinoma growth and aggressiveness via modulation of the ERK-JNK-mediated AKT/mTOR signaling pathway. Mol Med Rep 17, 4203–4212 (2018).

56. P. Lu, V. M. Weaver, Z. Werb, The extracellular matrix: A dynamic niche in cancer progression. Journal of Cell Biology 196, 395–406 (2012).

57. C. P. Anderson, M. Shen, R. S. Eisenstein, E. A. Leibold, Mammalian iron metabolism and its control by iron regulatory proteins. Biochimica et Biophysica Acta (BBA)-Molecular Cell Research 1823, 1468–1483 (2012).

58. M. U. Muckenthaler, S. Rivella, M. W. Hentze, B. Galy, A red carpet for iron metabolism. Cell 168, 344–361 (2017).

59. R. S. Eisenstein, K. P. Blemings, Iron regulatory proteins, iron responsive elements and iron homeostasis. J Nutr 128, 2295–2298 (1998).

60. M. Olea-Flores, M. Zuñiga-Eulogio, A. Tacuba-Saavedra, M. Bueno-Salgado, A. Sánchez-Carvajal, Y. Vargas-Santiago, M. A. Mendoza-Catalán, E. Pérez Salazar, A. García-Hernández, T. Padilla-Benavides, N. Navarro-Tito, Leptin Promotes Expression of EMT-Related Transcription Factors and Invasion in a Src and FAK-Dependent Pathway in MCF10A Mammary Epithelial Cells. Cells 8, 1133 (2019).

61. C. L. Roberge, L. Wang, M. Barroso, D. T. Corr, Non-Destructive Evaluation of Regional Cell Density Within Tumor Aggregates Following Drug Treatment. Journal of Visualized Experiments (JoVE) 184, e64030 (2022).

62. K. Tomita, M. Fukumoto, K. Itoh, Y. Kuwahara, K. Igarashi, T. Nagasawa, M. Suzuki, A. Kurimasa, T. Sato, MiR-7-5p is a key factor that controls radioresistance via intracellular Fe2+ content in clinically relevant radioresistant cells. Biochemical and Biophysical Research Communications 518, 712–718 (2019).

63. C. A. Schneider, W. S. Rasband, K. W. Eliceiri, NIH Image to ImageJ: 25 years of image analysis. Nature Methods 9, 671–675 (2012).

64. M. S. Messina, L. Torrente, A. T. Pezacki, H. I. Humpel, E. L. Li, S. G. Miller, O. Verdejo-Torres, T. Padilla-Benavides, D. C. Brady, D. W. Killilea, A. N. Killilea, M. Ralle, N. P. Ward, J. Ohata, G. M. DeNicola, C. J. Chang, A histochemical approach to activity-based copper sensing reveals cuproplasia-dependent vulnerabilities in cancer. Proceedings of the National Academy of Sciences 122, e2412816122 (2025).

65. O. Verdejo-Torres, D. C. Klein, L. Novoa-Aponte, J. Carrazco-Carrillo, D. Bonilla-Pinto, A. Rivera, A. Bakhshian, F. a. M. Fitisemanu, M. L. Jiménez-González, L. Flinn, Cysteine Rich Intestinal Protein 2 is a copper-responsive regulator of skeletal muscle differentiation and metal homeostasis. PLoS genetics 20, e1011495 (2024).

66. B. Alibayov, A. Scasny, F. Khan, A. Creel, P. Smith, A. G. J. Vidal, F. a. M. Fitisemanu, T. Padilla-Benavides, J. N. Weiser, J. E. Vidal, Oxidative reactions catalyzed by hydrogen peroxide produced by Streptococcus pneumoniae and other streptococci cause the release and degradation of heme from hemoglobin. Infection and Immunity 90, e00471–00422 (2022).

67. C. Tavera-Montañez, S. J. Hainer, D. Cangussu, S. J. Gordon, Y. Xiao, P. Reyes-Gutierrez, A. N. Imbalzano, J. G. Navea, T. G. Fazzio, T. Padilla-Benavides, The classic metal-sensing transcription factor MTF1 promotes myogenesis in response to copper. The FASEB Journal 33, 14556 (2019).

68. M. M. Bradford, A rapid and sensitive method for the quantitation of microgram quantities of protein utilizing the principle of protein-dye binding. Anal Biochem 72, 248–254 (1976).

69. J. M. Kelm, N. E. Timmins, C. J. Brown, M. Fussenegger, L. K. Nielsen, Method for generation of homogeneous multicellular tumor spheroids applicable to a wide variety of cell types. Biotechnology and Bioengineering 83, 173–180 (2003).

70. M. T. Santini, G. Rainaldi, Three-Dimensional Spheroid Model in Tumor Biology. Pathobiology 67, 148–157 (1999).

71. J. Schindelin, I. Arganda-Carreras, E. Frise, V. Kaynig, M. Longair, T. Pietzsch, S. Preibisch, C. Rueden, S. Saalfeld, B. Schmid, J. Y. Tinevez, D. J. White, V. Hartenstein, K. Eliceiri, P. Tomancak, A. Cardona, Fiji: an open-source platform for biological-image analysis. Nature methods 9, 676–682 (2012).

72. M. Marotta, G. Martino, Sensitive spectrophotometric method for the quantitative estimation of collagen. Anal Biochem 150, 86–90 (1985).

73. R. R. Lareu, D. I. Zeugolis, M. Abu-Rub, A. Pandit, M. Raghunath, Essential modification of the Sircol Collagen Assay for the accurate quantification of collagen content in complex protein solutions. Acta Biomater 6, 3146–3151 (2010).

74. A. S. Baez-Gonzalez, J. A. Carrazco-Carrillo, G. Figueroa-Gonzalez, L. I. Quintas-Granados, T. Padilla-Benavides, O. D. Reyes-Hernandez, Functional effect of indole-3 carbinol in the viability and invasive properties of cultured cancer cells. Biochem Biophys Rep 35, 101492 (2023).

75. M. L. Lacombe, F. Lamarche, O. De Wever, T. Padilla-Benavides, A. Carlson, I. Khan, A. Huna, S. Vacher, C. Calmel, C. Desbourdes, C. Cottet-Rousselle, I. Hininger-Favier, S. Attia, B. Nawrocki-Raby, J. Raingeaud, C. Machon, J. Guitton, M. Le Gall, G. Clary, C. Broussard, P. Chafey, P. Therond, D. Bernard, E. Fontaine, M. Tokarska-Schlattner, P. Steeg, I. Bieche, U. Schlattner, M. Boissan, The mitochondrially-localized nucleoside diphosphate kinase D (NME4) is a novel metastasis suppressor. BMC Biol 19, 228 (2021).

76. H. J. Bowers, E. E. Fannin, A. Thomas, J. A. Weis, Characterization of multicellular breast tumor spheroids using image data-driven biophysical mathematical modeling. Scientific Reports 10, 11583 (2020).

77. K. M. Charoen, B. Fallica, Y. L. Colson, M. H. Zaman, M. W. Grinstaff, Embedded multicellular spheroids as a biomimetic 3D cancer model for evaluating drug and drug-device combinations. Biomaterials 35, 2264–2271 (2014).

78. D. Loessner, K. S. Stok, M. P. Lutolf, D. W. Hutmacher, J. A. Clements, S. C. Rizzi, Bioengineered 3D platform to explore cell–ECM interactions and drug resistance of epithelial ovarian cancer cells. Biomaterials 31, 8494–8506 (2010).

79. M. Vinci, S. Gowan, F. Boxall, L. Patterson, M. Zimmermann, W. Court, C. Lomas, M. Mendiola, D. Hardisson, S. A. Eccles, Advances in establishment and analysis of three-dimensional tumor spheroid-based functional assays for target validation and drug evaluation. BMC Biology 10, 29 (2012).

80. M. Martin, Cutadapt removes adapter sequences from high-throughput sequencing reads. *EMBnet*. Journal 17, 10–12 (2011).

81. Y. Liao, G. K. Smyth, W. Shi, The R package Rsubread is easier, faster, cheaper and better for alignment and quantification of RNA sequencing reads. Nucleic Acids Res 47, e47 (2019).

82. Y. Liao, G. K. Smyth, W. Shi, featureCounts: an efficient general purpose program for assigning sequence reads to genomic features. Bioinformatics 30, 923–930 (2013).

83. C. W. Law, Y. Chen, W. Shi, G. K. Smyth, voom: Precision weights unlock linear model analysis tools for RNA-seq read counts. Genome Biol 15, R29 (2014).

84. M. E. Ritchie, B. Phipson, D. Wu, Y. Hu, C. W. Law, W. Shi, G. K. Smyth, limma powers differential expression analyses for RNA-sequencing and microarray studies. Nucleic Acids Research 43, e47–e47 (2015).

85. Y. Liao, J. Wang, E. J. Jaehnig, Z. Shi, B. Zhang, WebGestalt 2019: gene set analysis toolkit with revamped UIs and APIs. Nucleic Acids Res 47, W199–w205 (2019).

