## Supplementary material for "Iron–endoplasmic reticulum–extracellular matrix axis regulates cancer cell invasion"

1 Dept. Molecular and Cellular Physiology, Albany Medical College, Albany, NY 12208 USA

2 Molecular Biology and Biochemistry Department, Wesleyan University, Middletown, CT 06459 USA.

3 Dept. Biomedical Engineering, Rensselaer Polytechnic Institute, Troy, NY 12180 USA

4 Center for Modeling, Simulation, and Imaging in Medicine (CeMSIM), Rensselaer Polytechnic Institute, Troy, NY 12180 USA

5 Department of Research & Engineering, PixeSci, Inc., SUNY Albany, NY 12203 USA

Corresponding author: Margarida Barroso

### Material and Methods

**Optical coherence tomography imaging.** OCT imaging used a spectral-domain system (TEL220C1, Thorlabs) at a 1310 nm central wavelength, with axial resolution of 5.5 μm in air (4.2 μm in water) and lateral resolution of 5 μm (32). A-scans were collected at 5.5 kHz with 101 dB sensitivity, a medium refractive index of 1.33, and a 1.0 μm lateral pixel size. Volumes were exported to Imaris (v10.0; Oxford Instruments), where spheroid surfaces were reconstructed by manually tracing the boundary in three dimensions. Imaris then computed spheroid volume and sphericity, the latter defined as the surface area of an equivalent-volume ideal sphere divided by the measured surface area (1.0 = perfect sphere). Cell distribution was quantified with the Spot Finder module, detecting cells as 10 μm spots within the spheroid surface.

**Live cell imaging.** For live-cell imaging, No. 1.5 glass-bottom dishes (MatTek) were coated with poly-D-lysine to improve attachment. Cells were imaged in DHB medium (phenol red–free DMEM, 25 mM HEPES, L-glutamine, 0.5% BSA, pH 7.2) at 37 °C under 5% $CO_2$ on a Leica Thunder microscope fitted with a stage-top incubator. Fluorescence was recorded with filter sets matched to excitation/emission maxima of 488/509 nm (GFP) and 493/505 nm (ZsGreen). Images were analyzed in ImageJ or Imaris depending on the measurement.

**FerroOrange live-cell assay.** Labile iron pool (LIP) visualization was performed with FerroOrange staining (Dojindo Molecular Technologies) to detect labile ferrous iron ($Fe^{2+}$) (62). For 2D cultures, $1 \times 10^5$ cells were plated in complete DMEM on No. 1.5 poly-D-lysine-coated glass-bottom 35-mm dishes (MatTek) and cultured for 48 h. Before imaging, cells were washed twice with HBSS ($Ca^{2+}$/$Mg^{2+}$) and pre-incubated for 45 min at 37°C in DHB medium. A 1 μM working solution of FerroOrange was freshly prepared from stock under low light, and monolayers were stained for 15 min at 37°C. Cells were then rinsed three times with warm HBSS and kept in DHB medium during imaging. Spheroids were processed similarly with minor changes: 4-day spheroids were transferred to poly-D-lysine-coated glass-bottom dishes, washed in HBSS, and pre-incubated in DHB medium for 45 min at 37°C before staining with 1 μM FerroOrange for 15 min at 37 °C, washed three times in HBSS, and kept in DHB medium for live-cell imaging. 2D cultures were acquired with a 63× objective as Z-stacks (7 slices); spheroids were imaged with a 20× objective as a complete Z-stack through the structure. FerroOrange fluorescence was detected at 555 nm excitation and 595 nm emission on a Leica Thunder DMi8 microscope (Leica Microsystems) run through LAS X (Leica Microsystems). Mean FerroOrange intensity per cell or spheroid was measured in ImageJ (NIH) (63).

**Immunoblotting.** Monolayer cells were washed twice with ice-cold PBS. For spheroid samples, only those cultured in 5% (v/v) Matrigel for 4 days were used, as this low Matrigel concentration contains negligible collagen I and does not interfere with protein quantification. Spheroids were collected by gentle centrifugation (200 × g, 5 min, 4°C), washed three times in ice-cold PBS to remove residual Matrigel, and pelleted. Monolayer and spheroid samples were lysed on ice in 25 mM HEPES, 150 mM NaCl, 1 mM $MgCl_2$, and 0.4% NP-40 (pH 8) with protease and phosphatase inhibitors (Millipore), incubated 30 min with intermittent mixing, and clarified at 14,000 × g for 15 min at 4 °C. Protein was measured by BCA assay (Thermo Fisher). Equal protein (20-40 µg) was resolved by SDS-PAGE and transferred to PVDF membranes (Bio-Rad). Membranes were blocked 8-10 min at room temperature in EveryBlot buffer (Bio-Rad) and incubated overnight at 4°C with primary antibodies in blocking buffer. After three TBST washes, membranes were probed with HRP-conjugated secondary antibodies for 1 h at room temperature, with β-actin as loading control. Bands were visualized by enhanced chemiluminescence (ECL, Thermo Fisher) on a ChemiDoc system (Bio-Rad) and quantified by densitometry in ImageJ (NIH) (63).

**Metal content analysis.** Intracellular metal content was measured by atomic absorption spectroscopy (AAS) as described (33-35, 40, 41, 64-67). WT, KO, and OE cells were grown to 80-90% confluency, washed three times in ice-cold PBS (Gibco), scraped into 500 µl ultra-pure water (Thermo Fisher), and collected in a microcentrifuge tube. Lysates were sonicated (Bioruptor, medium setting, 30-s on/off cycles, 5 min) and total protein was measured by Bradford assay (68). Samples were then mineralized in 300 µl nitric acid (TraceMetal Grade, Fisher Chemical), boiled for 2 h, and left overnight at room temperature. They were neutralized with 10% $H_2O_2$ and brought to a final volume of 1.5 ml. Total iron was quantified on a 55B flame atomic absorption spectrometer (Agilent). To avoid trace contamination, all reagents and standards were analytical grade and prepared in 18 MΩ purified water, and glassware was soaked in 3% $HNO_3$ for 24 h before use. Iron standards (Thermo Fisher) were diluted in ultra-pure water to set the detection limit and dynamic range. Iron content was normalized to total protein.

**Second harmonic generation (SHG) imaging of collagen in spheroids.** WT, KO, and OE MDA-MB-231 cells were seeded in U-bottom, ultra-low attachment 96-well plates to form spheroids by liquid overlay (69, 70). After 4 days, mature spheroids were transferred to Ibidi 8-well µ-slides and left to settle for at least 30 min in phenol red-free medium before imaging. Collagen was visualized label-free by second harmonic generation on a multiphoton microscope with a 25× water-immersion objective. The SHG signal was excited at 820 nm and collected at 410 nm with a nondescanned detector. Z-stacks spanning the whole spheroid were acquired with identical laser power, detector gain, pixel size, and step size across conditions. Orthogonal (x-z and y-z) views were generated from the 3D stacks in the microscope software and exported as 8-bit grayscale TIFF files. Only linear adjustments were applied, identically to WT, KO, and OE spheroids. Imaging was performed on a Nikon AX MP multiphoton microscope (Nikon Instruments) with acquisition in NIS Elements (Nikon). Fluorescence intensity of SHG was quantified across 6 spheroids per condition in each experiment using ImageJ (63, 71).

**Sircol soluble collagen assay.** Soluble collagen secreted into conditioned medium was measured with the Sircol Soluble Collagen Assay (Biocolor, supplied through Ilex Life Sciences; Cat# S1000) according to the manufacturer's instructions, using the supplied bovine collagen standard (72). Cultures were maintained in phenol red-free DMEM to avoid absorbance interference. Medium was collected from 2D monolayers and day-4 LOL spheroids, cleared by centrifugation, and assayed against blanks of fresh medium (2D) or 5% Matrigel in medium (spheroids), so that reported values exclude collagen contributed by Matrigel. Absorbance was read at 556 nm and collagen normalized to 500,000 cells (n = 6 per condition). Because Sirius Red also binds non-collagenous proteins in serum-containing medium, values are compared between genotypes rather than treated as absolute collagen mass (73).

**Matrigel invasion assay in monolayer culture.** Transwell invasion assays were carried out as described (40, 41, 74) using BioCoat® Matrigel® Invasion Chambers with 8.0 μm PET membranes (Corning). WT, KO, and OE MDA-MB-231 cells were pretreated for 2 h with 10 μM cytosine β-D-arabinofuranoside (AraC) to suppress proliferation during the assay, then seeded at 125,000 cells/ml in 2 ml serum-free medium in the upper chamber. The lower chamber held 2.5 ml DMEM with 10% FBS as chemoattractant. After 72 h, non-invading cells and Matrigel were removed from the upper membrane with a cotton swab, and invading cells on the underside were washed, fixed in methanol for 5 min, and stained with 0.1% crystal violet in PBS. Six independent biological replicates were imaged with a 10× objective on an Echo Rebel microscope and quantified in FIJI v1.44p (71).

**Wound healing assay.** WT, KO, and OE cells were grown to confluency in 24-well plates in DMEM with 10% FBS and 1% penicillin/streptomycin. Medium was switched to serum-free for 24 h, after which cells were treated with 10 μM AraC for 2 h to arrest cell division during the assay. Monolayers were scratched with a sterile 200 μl pipette tip, and detached cells were rinsed away with PBS (40, 41, 74, 75). A reference mark on the plate allowed the same field to be imaged at the scratch (0 h) and at intervals thereafter until closure, using a 10× objective on an Echo Rebel microscope (Discover Echo). Images were analyzed in FIJI v1.44p (71).

**Three-dimensional invasion assay.** Spheroid invasion assays followed published protocols (76-79) with minor modifications. LOL spheroids were formed in CELLSTAR® Cell-Repellent 96-well U-bottom plates (VWR) with 5% Matrigel and matured for 4 days. To assay invasion into a denser matrix, mature spheroids were transferred to Cellvis 96-well glass-bottom plates (1.5 high-performance cover glass) and embedded in 20% Matrigel or 20% Type I Collagen in complete DMEM, mimicking a thicker ECM.

Spheroids were stimulated with 40 ng/ml TGF-β1, a known driver of breast cancer invasion (43). Plates were held at 37°C under 5% $CO_2$ in a humidified chamber, with perimeter wells filled with PBS to limit evaporation. Tumor cells were imaged by ZsGreen fluorescence on a Leica Thunder microscope at 5×, and Z-stacks spanning the spheroid and invasive front were collected on days 1, 4, and 8. To test the effect of iron depletion and ER stress on spheroid invasion, drug treatments were applied under two conditions. For iron chelation, spheroids received deferoxamine (DFO) at 5, 25, or 50 μM; for ER stress, tunicamycin at 1 or 5 μg/ml. All treatments were added at the time of embedding in 20% Matrigel, and invasion and morphology were followed by live imaging as above. Invasion was quantified in FIJI/ImageJ (63, 71). Z-stacks were converted to maximum-intensity projections and then to 32-bit grayscale, thresholded uniformly, and invasion was measured as the fluorescent area of the spheroid plus invasive protrusions. Invasion was scored as the increase in fluorescent area from day 4 to day 8, because spheroids showed little to no outward migration during the first 4 days as cells adapted to the surrounding matrix, and the change in fluorescent area between days 1 and 4 was negligible.

**RNA sequencing and bioinformatics analysis.** RNA quality was checked on an Agilent Bioanalyzer, and all samples had RNA Integrity Numbers above 8.6. Libraries were prepared by poly(A) selection and sequenced on an Illumina HiSeq (2×150 bp paired-end) by GENEWIZ. Raw FASTQ files were assessed with FastQC and trimmed for adaptors and quality with Trim Galore (80). Reads were aligned to the human genome (hg38) with Rsubread v1.5.3 (81), and gene-level counts were obtained with featureCounts (82), annotated by Entrez Gene IDs. Genes with counts-per-million > 0.5 in at least three samples were retained. Differential expression was assessed with the limma-voom pipeline (83, 84), with significance set at an adjusted p-value < 0.05. Functional enrichment was performed with WebGestalt (85).

Transcriptomic data (Fig. 6A) were drawn from the METABRIC breast cancer cohort via cBioPortal (Curtis et al.; Pereira et al.), restricted to basal-like tumors with complete expression values for all 10 panel markers and one sample per patient ($n = 198$). Proteomic data (Fig. 6B) came from the CPTAC cohort through the NCI Proteomic Data Commons, retaining samples with complete quantitation for the same 10 markers ($n = 55$).

The same selected 10-marker panel was applied to both datasets, spanning iron regulation (SLC11A2/DMT1, FTH1), ER stress and proteostasis (HSPA5, P4HB, ERO1A, P4HA1), and ECM/EMT programs (ITGB1, ICAM1, TGFB1, HIF1A). Pairwise associations were computed as Spearman rank coefficients (ρ) with two-sided tests within each cohort; significance markers in Fig. 6 reflect nominal p-value thresholds (*$p < 0.05$, **$p < 0.01$, ***$p < 0.001$). To compare cohorts, the 45 shared off-diagonal coefficients from the

METABRIC and CPTAC matrices were extracted and related by Pearson and Spearman correlation.

Correlation matrices were built in Python 3.12 (numpy, pandas, SciPy, seaborn, matplotlib, statsmodels), hierarchically clustered to group related markers, and annotated with pathway-category side bars for the iron, ER stress, and ECM/EMT sets.

**Supplementary Figures**

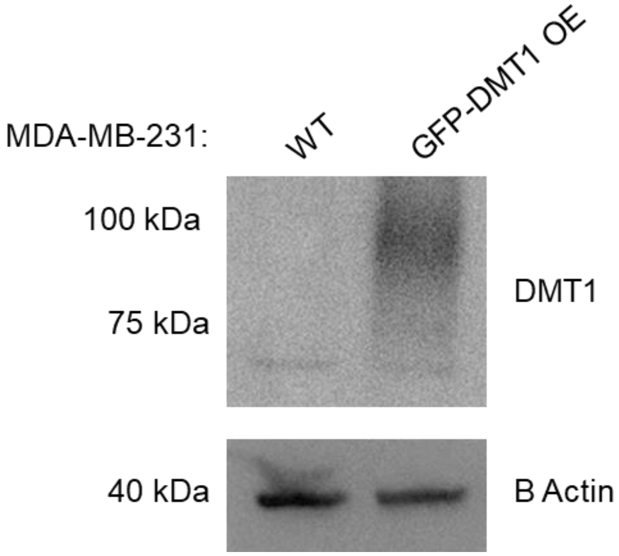

**Fig. S1. Validation of stable DMT1-GFP overexpression in MDA-MB-231 cells**. Western blot analysis confirmed stable expression of the DMT1-GFP overexpression construct in MDA-MB-231 OE cells before downstream transcriptomic and phenotypic analyses. OE cells showed elevated DMT1-GFP fusion protein signal relative to WT controls. β-actin served as a loading control.

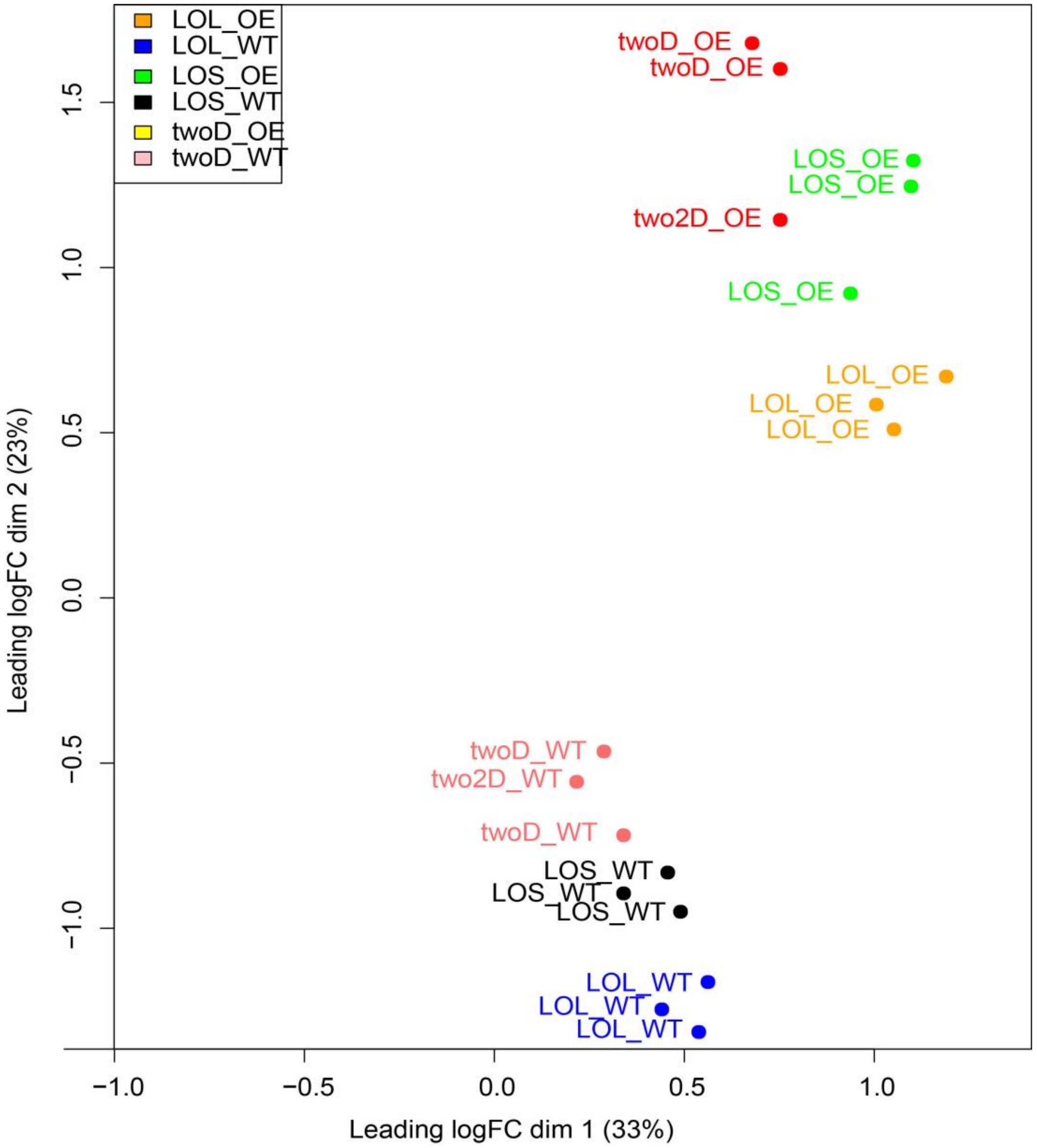

**Fig. S2. Principal component analysis (PCA) of transcriptomic profiles across 2D, LOS, and LOL cultures in wild-type (WT) and DMT1-GFP overexpressing (OE) MDA-MB-231 cells.** PCA of RNA sequencing data shows that culture dimensionality and spheroid size were the dominant drivers of global transcriptional variance, whereas DMT1 overexpression produced comparatively smaller within-condition shifts. PC1 (33% variance) primarily separated 2D cultures from 3D spheroids, with LOL constructs, showing the greatest displacement from 2D conditions, indicating that larger spheroids induce the most substantial transcriptional remodeling. PC2 (23% variance) showed only minor separation between WT and OE cells within each culture condition. LOS and LOL samples formed distinct clusters, confirming progressive size-dependent physiological divergence, while OE samples largely preserved the same dimensional trajectory as WT counterparts. These findings indicate that DMT1 overexpression does not override the size-dependent transcriptional baseline of the cell culture model.

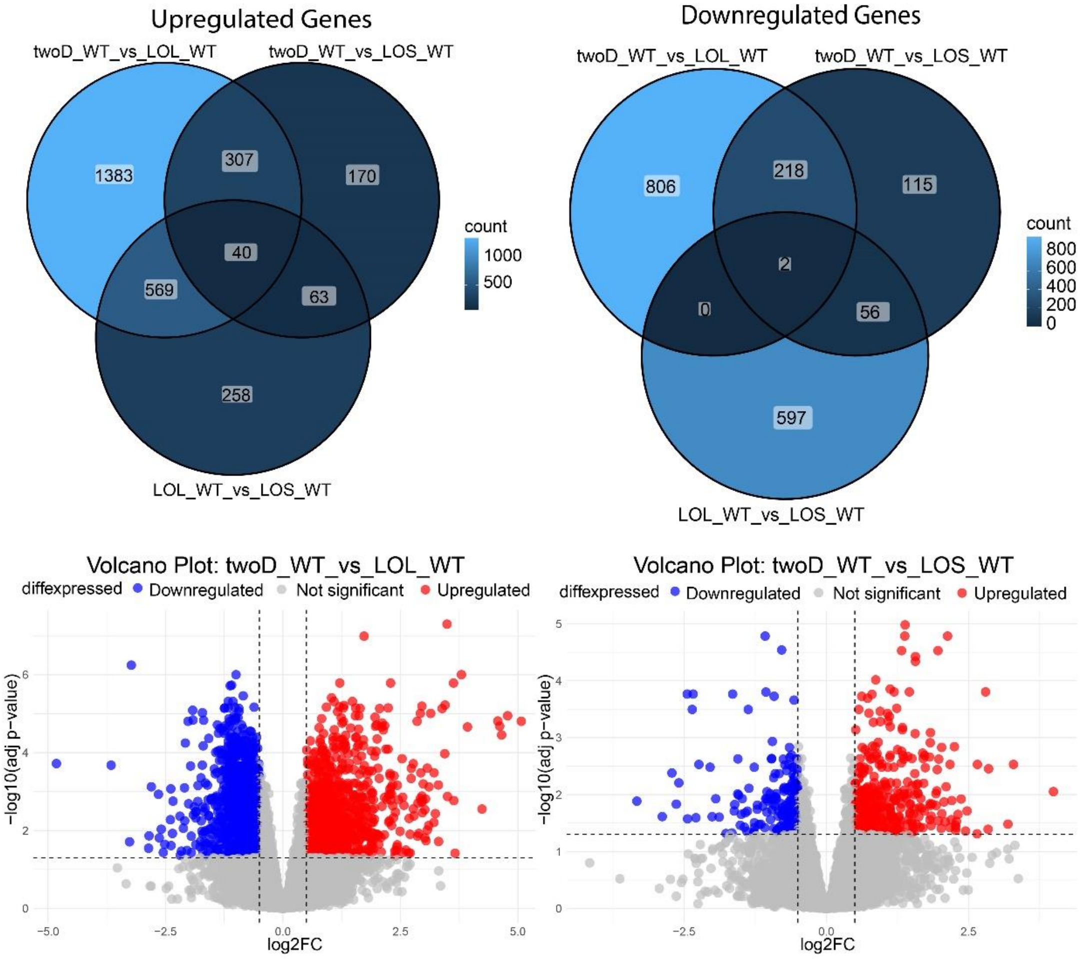

**Fig. S3. Differential gene expression in WT MDA-MB-231 cells across 2D, LOL, and LOS conditions. (A)** Venn diagrams showing the overlap of upregulated genes among the WT MDA-MB-231 comparisons of 2D vs. LOL, 2D vs. LOS, and LOL vs. LOS. **(B)** Venn diagrams showing the overlap of downregulated genes for the same comparisons. **(C)** Volcano plot for 2D WT vs. LOL WT showing significantly upregulated (red) and downregulated (blue) transcripts based on log2 fold-change and adjusted p-value thresholds. **(D)** Volcano plot for 2D WT vs. LOS WT showing the distribution of differentially expressed genes under the same criteria. The complete set of differentially expressed genes across these comparisons is provided in Data file S1.

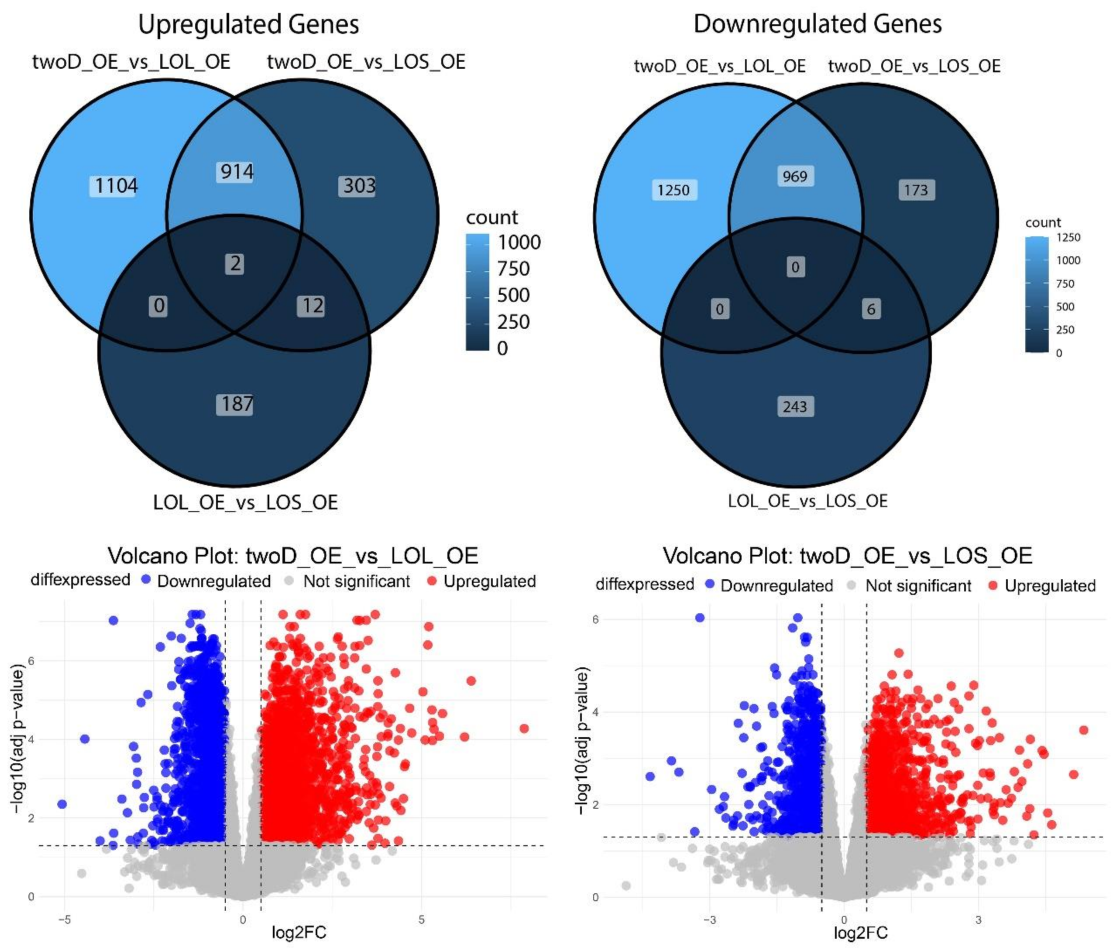

**Fig. S4. Differential gene expression in DMT1-overexpressing (OE) cells across 2D, LOL, and LOS conditions. (A)** Venn diagrams showing the overlap of upregulated genes among the OE MDA-MB-231 comparisons of 2D vs. LOL, 2D vs. LOS, and LOL vs. LOS. **(B)** Venn diagrams showing the overlap of downregulated genes for the same three comparisons. **(C)** Volcano plot for 2D OE vs. LOL OE highlighting significantly upregulated (red) and downregulated (blue) transcripts based on log2 fold-change and adjusted p-value thresholds. **(D)** Volcano plot for 2D OE vs. LOS OE showing the distribution of differentially expressed genes under the same significance criteria. The complete set of differentially expressed genes across these comparisons is provided in Data file S3.

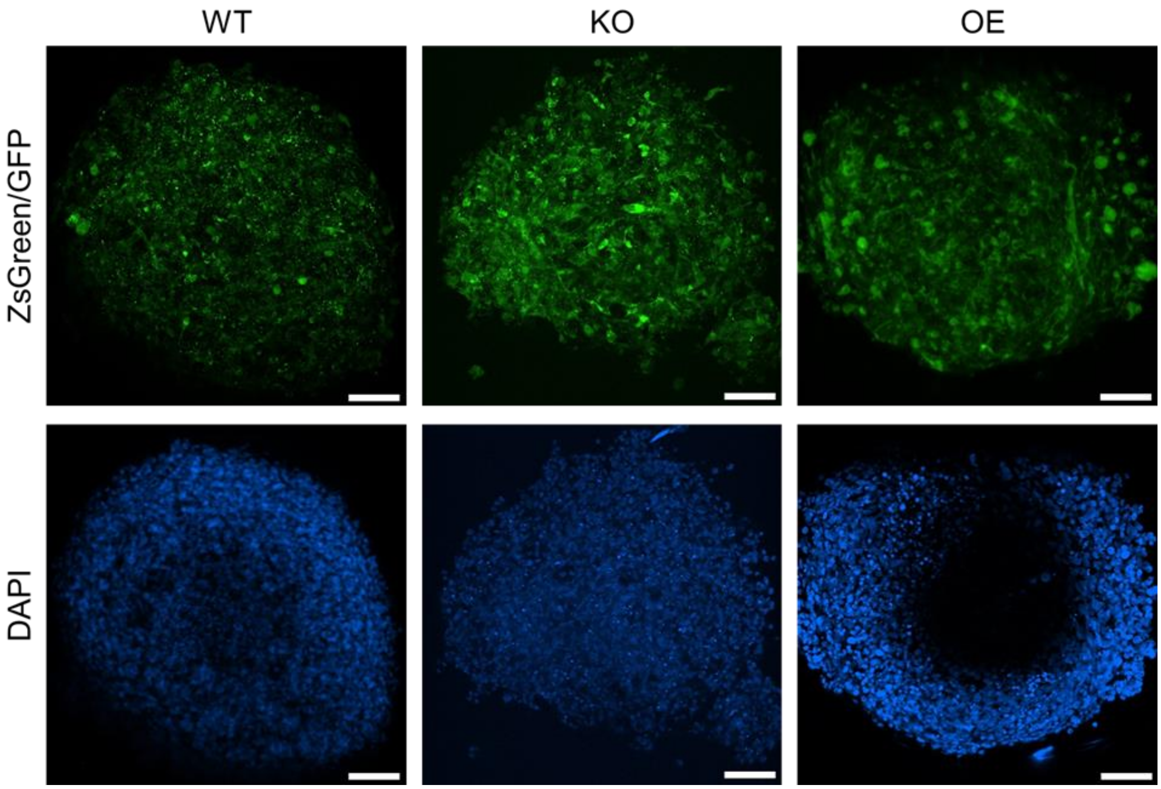

**Fig. S5. Two-photon microscopy of MDA-MB-231 spheroids with altered DMT1 expression.** Maximum-intensity projection images of freshly fixed MDA-MB-231 LOL spheroids expressing ZsGreen/GFP (top row) and stained with DAPI for 10 minutes (bottom row). WT and KO spheroids were visualized by ZsGreen fluorescence, whereas OE spheroids were visualized by DMT1-GFP fluorescence**.** WT, DMT1 KO, and OE spheroids were imaged using a 25× objective. In OE spheroids, the dense and compact core restricted DAPI penetration, resulting in weak nuclear staining despite uniform GFP fluorescence throughout the spheroid. WT and KO spheroids showed more uniform DAPI labeling, consistent with less restricted dye diffusion. Scale bar, 100 µm.

### 1. Tables

**Table S1.** Key resources table.

Comprehensive list of reagents, antibodies, cell lines, deposited datasets, and software used in this study, including source vendors and catalog numbers or accession identifiers. This table provides full resource transparency for experimental reproducibility across biochemical assays, 3D spheroid culture, imaging, RNA sequencing analyses, and computational workflows.

| REAGENT or RESOURCE | SOURCE | IDENTIFIER |
|---|---|---|
| **Chemicals, peptides and recombinant proteins** | | |
| Protease inhibitor cocktail | Roche | Cat# 04693116001 |
| 4X Laemmli sample buffer | BioRad | Cat# 1610747 |
| 4-20% Mini-Protean TGX gels | BioRad | Cat# 4561094 |
| Bovine Serum Albumin | Millipore-Sigma | Cat# A9647 |
| 2-b-Mercaptoethanol | Millipore-Sigma | Cat# M3148 |
| Clarity Max Western ECL | BioRad | Cat# 1705062 |
| DMEM high glucose, no glutamine, no phenol red | Thermo Fisher Scientific | Cat# 31053-028 |
| Fetal Bovine serum (FBS) | ATCC | Cat# 30-2020 |
| DAPI | Thermo Fisher Scientific | Cat# D1306 |
| FerroOrange | Dojindo | Cat# F374 |
| Matrigel® Matrix | BD Biosciences | Cat# 356237 |
| Type I Collagen | ibidi | 50201 |
| Tunicamycin | Fisherscientific | 35-161-0 |
| TGF-B1 | Thermofisher | PHG9214 |
| DFO | Fischer Scientific | AC461770010 |

| | | |
|---|---|---|
| **Antibodies** | | |
| Rabbit anti-FTH1 | Cell Signaling | Cat# 3998 |
| Rabbit anti-ZIP8 | Cell Signaling | Cat# 47188S |
| Rabbit anti-IRP2 | Cell Signaling | Cat# 37135S |
| Rabbit anti-BIP | Cell Signaling | Cat# 3177 |
| Rabbit anti-ATF-6 | Cell Signaling | Cat# 65880 |
| Rabbit anti-IRE1 | Cell Signaling | Cat# 3294 |
| Rabbit anti-ERO1 | Cell Signaling | Cat# 3264 |
| Rabbit anti-PDI | Cell Signaling | Cat# 3501S |
| Rabbit anti-P4HA1 | Cell Signaling | Cat# 61182S |
| Rabbit anti-COL1A | ThermoFisher | PA1-26204 |
| Rabbit anti-ITGB1 | Cell Signaling | Cat# 9699S |
| Rabbit anti-TGFB1 | Cell Signaling | Cat# 3711S |
| Rabbit anti-ICAM1 | Cell Signaling | Cat# 67836S |
| Rabbit HRP-conjugated IgG | Cell Signaling | Cat# 7074S |
| Mouse HRP-conjugated IgG | Cell Signaling | Cat# 7076S |
| **Experimental models: Cell lines** | | |
| MDA-MB-231 | ATCC | Cat# HTB-26 |
| **Deposited data** | | |
| RNAseq of MDA-MB-231 WT vs DMT1 OE | This paper | GSE312425 |

| | | |
|---|---|---|
| RNAseq of MDA-MB-231 WT vs DMT1 KO | Bara et al, 2024 | GSE226059 |
| **Software and algorithms** | | |
| Imaris software | Oxford Instruments | https://imaris.oxinst.com/ |
| FIJI | NIH | https://imagej.net/software/fiji/ |
| Graphpad Prism 10.0 | Prism | https://www.graphpad.com |
| R package | The R Project for Statistical Computing | https://www.r-project.org/ |
| Python (v3.x) with pandas, NumPy, SciPy, scikit-learn, matplotlib, seaborn in Kaggle Notebooks | Python Software Foundation; Kaggle/Google LLC | https://www.python.org/ ; https://www.kaggle.com/ |
| LASX software | Leica | https://www.leica-microsystems.com/products/microscope-software/p/leica-las-x-ls |

**Table S2.** Source of Experimental Datasets Used in This Study.

| Experimental condition | Cell line | Genotype | Model | Generated by | GEO | Reference |
|---|---|---|---|---|---|---|
| 2D | MDA-MB-231 | Wild type (WT) | 2D Monolayer | Arun Asif | GSE312425 | This study |
| 2D | MDA-MB-231 | DMT1 Overexpression (OE) | 2D Monolayer | Arun Asif | GSE312425 | This study |
| LOS | MDA-MB-231 | Wild type (WT) | 3D spheroid Liquid-overlay small (LOS) | Arun Asif | GSE312425 | This study |

| | | | | | | |
|---|---|---|---|---|---|---|
| LOS | MDA-MB-231 | DMT1 Overexpression (OE) | 3D spheroid Liquid-overlay small (LOS) | Arun Asif | GSE312425 | This study |
| LOL | MDA-MB-231 | Wild type (WT) | 3D spheroid Liquid-overlay large (LOL) | Arun Asif | GSE312425 | This study |
| LOL | MDA-MB-231 | DMT1 Overexpression (OE) | 3D spheroid Liquid-overlay large (LOL) | Arun Asif | GSE312425 | This study |
| 2D | MDA-MB-231 | Wild type (WT) | 2D Monolayer | Jonathan Barra | GSE226059 | Barra et al., Oncogene 2024 (*17*) |
| 2D | MDA-MB-231 | DMT1 Knockout (KO) | 2D Monolayer | Jonathan Barra | GSE226059 | Barra et al., Oncogene 2024 (*17*) |
| LOL | MDA-MB-231 | Wild type (WT) | 3D spheroid Liquid-overlay large (LOL) | Jonathan Barra | GSE226059 | Barra et al., Oncogene 2024 (*17*) |
| LOL | MDA-MB-231 | DMT1 Knockout (KO) | 3D spheroid Liquid-overlay large (LOL) | Jonathan Barra | GSE226059 | Barra et al., Oncogene 2024 (*17*) |
| Patient cohort | Breast cancer (METABRIC) | Basal-like / TNBC subset | Human primary tumor transcriptome dataset | cBioPortal / METABRIC Consortium | METABRIC | Pereira et al., Nat Commun 2016; Curtis et al., Nature 2012 |

| | | | | | | |
|---|---|---|---|---|---|---|
| Patient cohort | Breast cancer (METABRIC) | All molecular subtypes | Human primary tumor transcriptome dataset | cBioPortal / METABRIC Consortium | METABRIC | Pereira et al., Nat Commun 2016; Curtis et al., Nature 2012 |
| Patient cohort | Breast cancer (CPTAC) | Breast invasive carcinoma proteome | Human primary tumor proteogenomic dataset | CPTAC Consortium | CPTAC Breast Cancer | Krug et al., Cell 2020 |
| Patient cohort | Breast cancer (CPTAC) | Basal-like / TNBC subset | Human primary tumor proteogenomic dataset | CPTAC Consortium | CPTAC Breast Cancer | Krug et al., Cell 2020 |

**Movie S1 (separate file).** MDA-MB-231 WT LOL spheroid 2-photon SHG imaging of collagen Z-stack animation

**Movie S2 (separate file).** MDA-MB-231 *DMT1* KO LOL spheroid 2-photon SHG imaging of collagen Z-stack animation

**Movie S3 (separate file).** MDA-MB-231 *DMT1* OE LOL spheroid 2-photon SHG imaging of collagen Z-stack animation
